# Discovery of a distinct BAM complex in the Bacteroidetes

**DOI:** 10.1101/2025.01.31.636011

**Authors:** Augustinas Silale, Mariusz Madej, Katarzyna Mikruta, Andrew M. Frey, Adam J. Hart, Arnaud Baslé, Carsten Scavenius, Jan J. Enghild, Matthias Trost, Robert P. Hirt, Bert van den Berg

## Abstract

The BAM (β-barrel assembly machinery) complex is an evolutionarily conserved, multiprotein machine that catalyses the folding and membrane insertion of newly synthesised β-barrel outer membrane (OM) proteins in Gram-negative bacteria. Based on Proteobacteria, bacterial BAM is also structurally conserved, with an essential BamAD core and up to three auxiliary periplasmic lipoproteins of poorly defined function. Here we show, using structural biology, quantitative proteomics and functional assays, that the BAM complex is radically different within the Bacteroidetes, a large and important phylum widely distributed within the environment and animal microbiomes. Cryogenic electron microscopy (cryo-EM) structures of BAM complexes from the human gut symbiont *Bacteroides thetaiotaomicron* and the human oral pathogen *Porphyromonas gingivalis* show similar, seven-component complexes of ∼325 kDa in size with most of the mass in the extracellular space. In addition to canonical BamA and BamD, the complexes contain an integral OM protein named BamF that is essential and intimately associated with BamA, as well as four surface-exposed lipoproteins (SLPs) named BamG-J. Together, BamF-J form a large, extracellular dome that likely serves as an assembly cage for the β-barrel-SLP complexes that are a hallmark of the Bacteroidetes. Our data suggest that BAM functionality in Bacteroidetes is substantially expanded from that in Proteobacteria and underscores the importance of studying other phyla for a more complete understanding of fundamental biological processes.

## Introduction

The BAM (β-barrel assembly machinery) complex is a conserved, multiprotein machine that mediates the folding and membrane insertion of newly synthesised β-barrel outer membrane (OM) proteins. BAM complex structure and mechanism of action have been studied in detail in γ-Proteobacteria, particularly *E. coli*. These studies have shown that the archetypal BAM complex is ∼200 kDa in size and consists of an essential, 16-stranded integral β-barrel OM protein (OMP) named BamA, which has five periplasmic POTRA (polypeptide-transport-associated) domains that are decorated in the periplasmic space by the four lipoproteins BamB-E^1–6^. BamA is an atypical β-barrel in which the first and last β-strands can separate to form a lateral gate that is open towards the OM. Recent studies demonstrate that the “uncovered” first BamA strand acts as a template to fold nascent OMPs via a “hybrid-barrel budding” mechanism^7–9^, likely facilitated by local OM destabilisation/thinning^10^. The BamB-E lipoproteins likely increase the efficiency of this process in ways that are not yet clear despite two decades of study^11,12^. Except for BamD, they are dispensable for cell viability in the absence of suppressor mutations^13^. Delivery of unfolded substrates to the BAM complex is mediated by several periplasmic chaperones like SurA and Skp, which function in distinct ways^14–17^. Phylogenetic analysis suggests only minor variations for BAM complex composition across bacterial phyla, caused by variable presence of the non-essential components BamB, BamC and BamE^18^. In addition to β-barrel integral OMPs, the OM contains lipoproteins. In Proteobacteria, most of these are targeted to and inserted into the inner leaflet of the OM via the Lol pathway, resulting in periplasmic exposure^19^. In addition to these “classical” lipoproteins, surface lipoproteins (SLPs) also exist which, after being targeted to the OM, need to be flipped for surface exposure^20^. Structurally and mechanistically, little is known about SLP flipping, although the Proteobacterial OMP Slam (surface lipoprotein assembly modulator) is known to be involved in a BAM-independent fashion^21,22^. Importantly, the numbers of Proteobacterial SLPs are generally low (∼10 in *E. coli,* which lacks SLAM), and complexes between SLPs and β-barrel OMPs, exemplified by the TbpAB iron piracy system of pathogenic *Neisseria*^23^, are uncommon. By contrast, stable SLP-barrel complexes are ubiquitous and abundant within the Bacteroidetes, a large Gram-negative bacterial phylum that is widely distributed within the environment (soil, sea water) and in animal microbiomes. The model human gut symbiont *Bacteroides thetaiotaomicron* (*B. theta*) has an estimated 400-450 SLPs, based on the presence of the Bacteroidetes lipoprotein export signal (LES)^24,25^ in a large subset of *B. theta* lipoproteins. Many of these SLPs are present in complexes with large TonB-dependent transporters (TBDTs) that mediate uptake of a wide range of dietary and host glycans, vitamin B_12_ and iron-siderophores^26,27^. These SLP-barrel complexes are likely to be key for the success of the genus *Bacteroides* in the human gut, but how the SLPs are flipped across the OM and how the SLP-barrel complexes are subsequently assembled is completely unclear.

Here, we report, based on the hypothesis that SLP flipping and SLP-barrel assembly may be physically linked, the discovery of a distinct BAM complex that is widespread within the Bacteroidetes. Cryogenic electron microscopy (cryo-EM) structures of the BAM complexes from *B. theta* and the oral pathogen *Porphyromonas gingivalis* are similar and reveal a 7-protein, ∼325 kDa complex with most of its mass on the extracellular side. Remarkably, besides BamA and BamD subunits, Bacteroidetal BAM contains a 14-stranded integral OMP that is intimately associated with BamA. We name this subunit BamF, as it is a homologue of the *E. coli* long-chain fatty acid transporter FadL. Repeated failure to delete BamF-encoding genes both in *B. theta* and in *P. gingivalis* suggests it is essential. The remaining four components, named BamG-I, are all SLPs and form an unprecedented extracellular dome-like structure over BamA that we propose functions as a folding cage for newly assembled SLP-barrel complexes that are a hallmark of the Bacteroidetes phylum. Our data suggest that the functionality of the BAM complex in Bacteroidetes is expanded and substantially different from that in Proteobacteria and underscores the importance of moving beyond Proteobacteria to gain a more complete understanding of fundamental biological processes.

## Results

### BtBamA co-purifies with several OMPs of unknown function

We reasoned, based on the need to assemble large numbers of structurally divergent SLP-barrel complexes in *B. theta*, that a putative SLP flippase might be associated with the BAM complex. To test this hypothesis, we tagged the chromosomal copy of the gene encoding BtBamA (*bt3725*) with an N-terminal His_7_ tag and performed pulldowns from *B. theta* membrane extracts in dodecyl-maltoside (DDM). Several proteins co-purified with BtBamA_his_ and their identities were established via peptide fingerprinting of pooled SEC peak fractions (Fig. 1a) and quantitative proteomics on IMAC elutions (Fig. 1b). Surprisingly, besides BamD (Bt0573) and SurA (Bt3848) that constitute a “minimal” Proteobacterial BAM system, BamA_his_ also copurifies with an integral β-barrel OMP (Bt4367), two peptidyl-prolyl isomerases (PPIs; Bt3612 and Bt3949), and two lipoproteins of unknown function, Bt3727 and Bt4306. Inspection of the sequences shows that both PPIs and the other two lipoproteins have at least two acidic residues within the first six residues after the lipid anchor cysteine, suggesting they are SLPs^24^. The pulldown data support bioinformatics analyses that suggest Bacteroidetes lack homologues of BamC and BamE. Whether any Bacteroidetes have BamB is not clear, but our data strongly suggest that *B. theta* does not. AlphaFold^28^ predictions of the five proteins (ED Fig. 1a) did not give obvious clues about a function within a putative BAM complex, and none of the proteins have predicted structures consistent with that expected for an OM SLP flippase (*i.e*., a large barrel that can open laterally). Interestingly, Bt4367 is one of the four *B. theta* paralogs of the long-chain fatty acid transporter FadL from *E. coli*^29,30^, and structurally similar to Bt1785 (ED Fig. 1a,b). Bt4367 (FadL/Toluene_X) and the PPI Bt3612 (FKBP_C) are widespread (ED Fig. 2), whereas Bt3727 (DUF6242), Bt4306 (DUF4270) and the PPI Bt3949 (DUF4827) appear to be confined to the Bacteroidetes. Of these, Bt4306 is widespread, while Bt3727 and Bt3949 largely restricted to the Bacteroidia class (ED Fig. 2). *B. theta* has one paralog of Bt4306, Bt1786, which is ∼100 residues shorter than Bt4306 but has a similar predicted structure (ED Fig. 1b).

**Figure 1.**
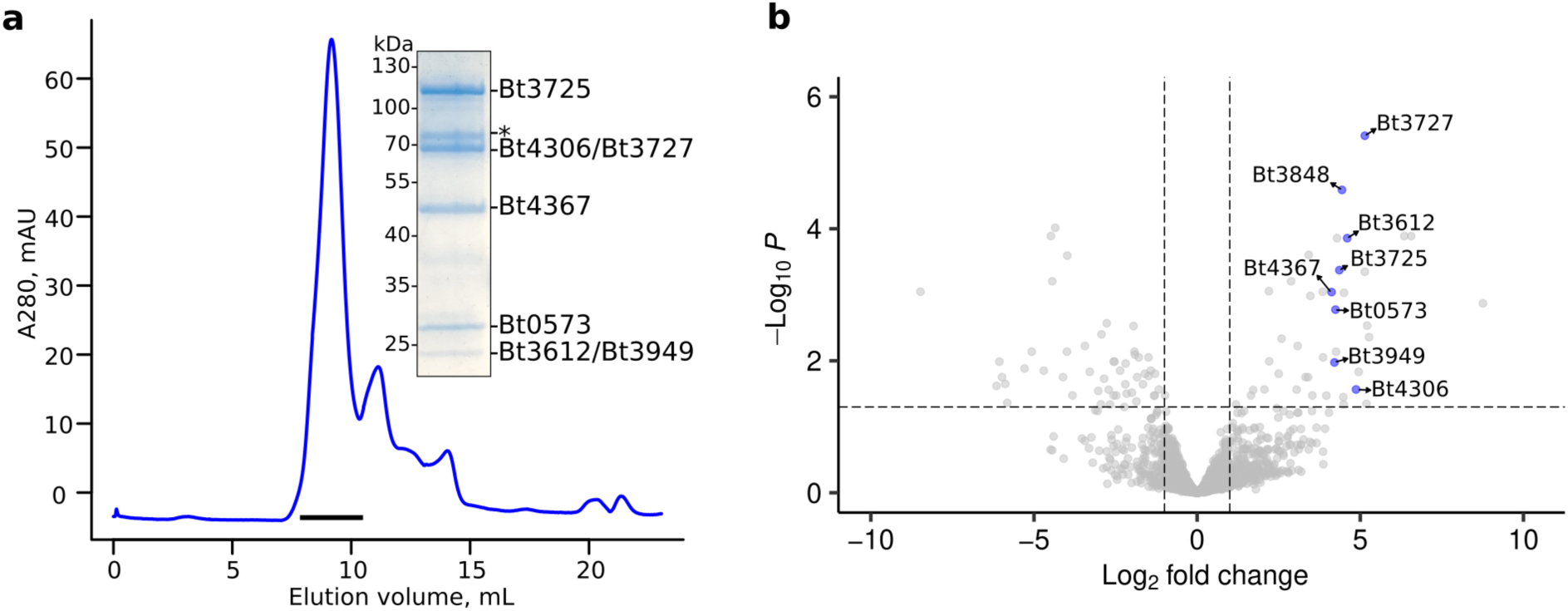
Identification of BtBAM components. **a**, Representative size exclusion chromatography trace of a BtBamA_his_ pulldown (n=5). Fractions denoted by the black horizontal bar were pooled, concentrated and analysed by 12% SDS-PAGE (inset). Individual bands were cut out and identified by peptide fingerprinting. The band marked with an asterisk corresponds to Bt0502. **b**, Volcano plot showing enrichment of proteins in pulldowns from *B. theta bamA_his_* versus wild type cells (n=3 biological replicates for both conditions). The vertical dashed lines represent log2 fold change=±1. Proteins above the horizontal dashed line have an adjusted p-value<0.05.

**Figure 2.**
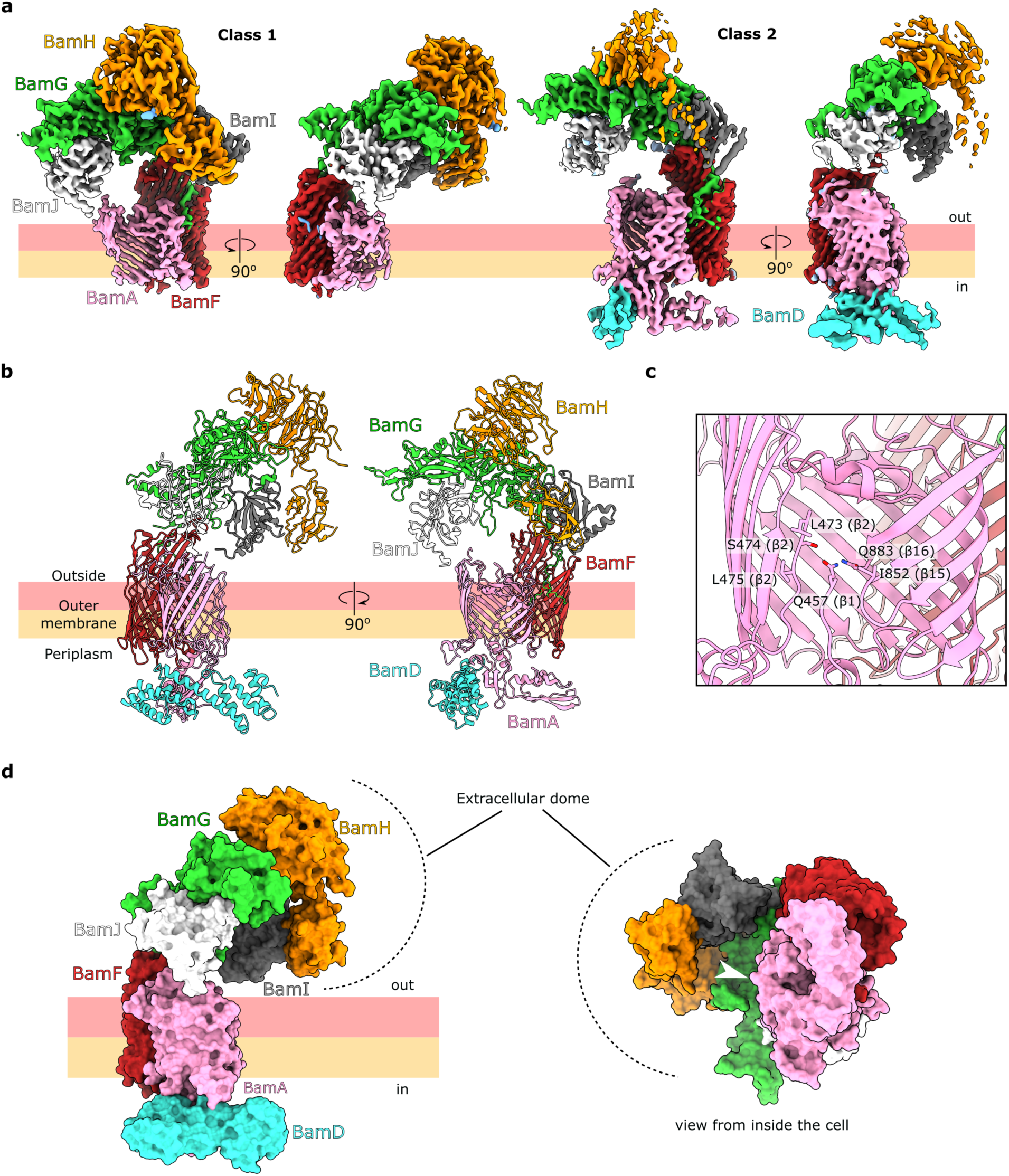
Single particle cryo-EM structure of BtBAM. **a**, Refined cryo-EM density maps of the final two 3D classes. **b**, Model built into the cryo-EM maps. The SLP models were refined against the class 1 density where the extracellular region was better resolved. The BamADF models were refined against class 2, where both the BamA and BamD density was better resolved. The refined SLP models were rigid-body-docked into the class 2 map, resulting in the final model. **c**, Close-up view of the BamA β-barrel seam. Residues 458-472, comprising most of the β1 strand, extracellular loop 1, and part of the β2 strand are not resolved in the cryo-EM maps. **d**, The SLPs form an extracellular dome, which partially encloses a ∼90,000 Å^3^ volume above the OM. The white arrowhead in the right panel indicates the lateral opening of the BamA β-barrel, with BamD omitted for clarity.

### The bulk of the BtBAM complex is extracellular

To obtain more information on the relationship of the copurified proteins with BtBamA, we collected single particle cryo-EM data on a sample in DDM purified from ∼10 liters of *B. theta* grown in rich medium. Maps to ∼3.3 Å resolution were obtained (Supplementary Fig. 1a-d; Methods) that allowed confident docking of AF2 models and subsequent refinement (Fig. 2a,b and Supplementary Table 1). All the abundant (*i.e*., visible as a clear band in SDS-PAGE after SEC) co-purified proteins except Bt0502 (Fig. 1a) are present in the cryo-EM volumes, each with one copy to generate a large complex of ∼325 kDa, which we term BtBAM (Fig. 2a,b). In striking contrast to Proteobacterial BAM, most of the BtBAM complex is extracellular due to the four newly discovered SLPs. Two particle classes are present in the dataset (Fig. 2a and Supplementary Fig. 1a). Class 1 lacks density for the periplasmic components but has good density for all extracellular proteins, whereas class 2 has modest resolution for BamA POTRA domains 4 and 5 and for part of BamD, but with poorly resolved extracellular areas (Supplementary Fig. 1e,f). Given that models derived from buildable areas in both maps are very similar with only small (max. ∼1 Å for Cα atoms) rigid body movements of the extracellular domain, we present only one, “hybrid” structural model. The dataset does not contain any particle populations of sub-complexes.

The 14-stranded barrel of the integral membrane component Bt4367 occupies a key position within BtBAM. It forms a tight complex with the back of BamA (defined as being located opposite the BamA lateral gate), with an interface area of ∼1560 Å^2^ as analysed via PISA^31^, and binds Bt4306 via its extracellular loops (interface area ∼1720 Å^2^) (Fig. 2b). The latter interaction is somewhat reminiscent to the proposed role of PorV, also a FadL homologue, as a substrate binding/shuttling protein in the Type IX secretion system (T9SS)^32^. However, given the intimate association of Bt4367 with both BamA and Bt4306, we consider it unlikely that Bt4367 has a PorV-like function in BtBAM. Because Bt4367 is a member of the FadL OMP family, we have named this subunit BamF. A feature that distinguishes BamF from FadL channels with a known transport role is an extension of the N-terminus of ∼10 residues, which protrudes through the lateral opening in the barrel wall that normally is part of the FadL transport pathway (ED Fig. 3a)^33,34^. Interestingly, a ConSurf^35^ analysis shows that the extended BamF N-terminus is highly conserved, but the extracellular loops interacting with Bt4306 are not (ED Fig. 3b). The extended N-terminus is also present in Bt1785 (ED Fig. 3a), but a detailed phylogenetic analysis shows that it, and the additional BamF encoded by *Bacteroides* spp. with more than one *bamF* gene, are distinct from Bt4367/BamF proteins (ED Fig. 3c,d). BlastP searches against the 1448 complete and annotated Bacteroidetes genomes in the RefSeq database (NCBI) using Bt4367/BamF as query identified one to three significant hits per genome in all but one free-living Bacteroidetes (Supplementary Table 2 and Supplementary Discussion). No significant hits were found in any endosymbiont Bacteroidetes. This is consistent with an important role played by BamF and its functional partners in processing cell surface proteins, the repertoire of which is typically dramatically reduced within the small genomes of endosymbiotic bacteria, which only maintain basic functionalities such as the biosynthesis of essential amino acids^36^. The other BtBAM components have been named BamG-I based on their distance to BamF, with Bt4306 corresponding to BamG, Bt3727 corresponding to BamH, and the PPIs Bt3612 and Bt3949 named BamI and BamJ, respectively (Fig. 2a,b).

**Figure 3.**
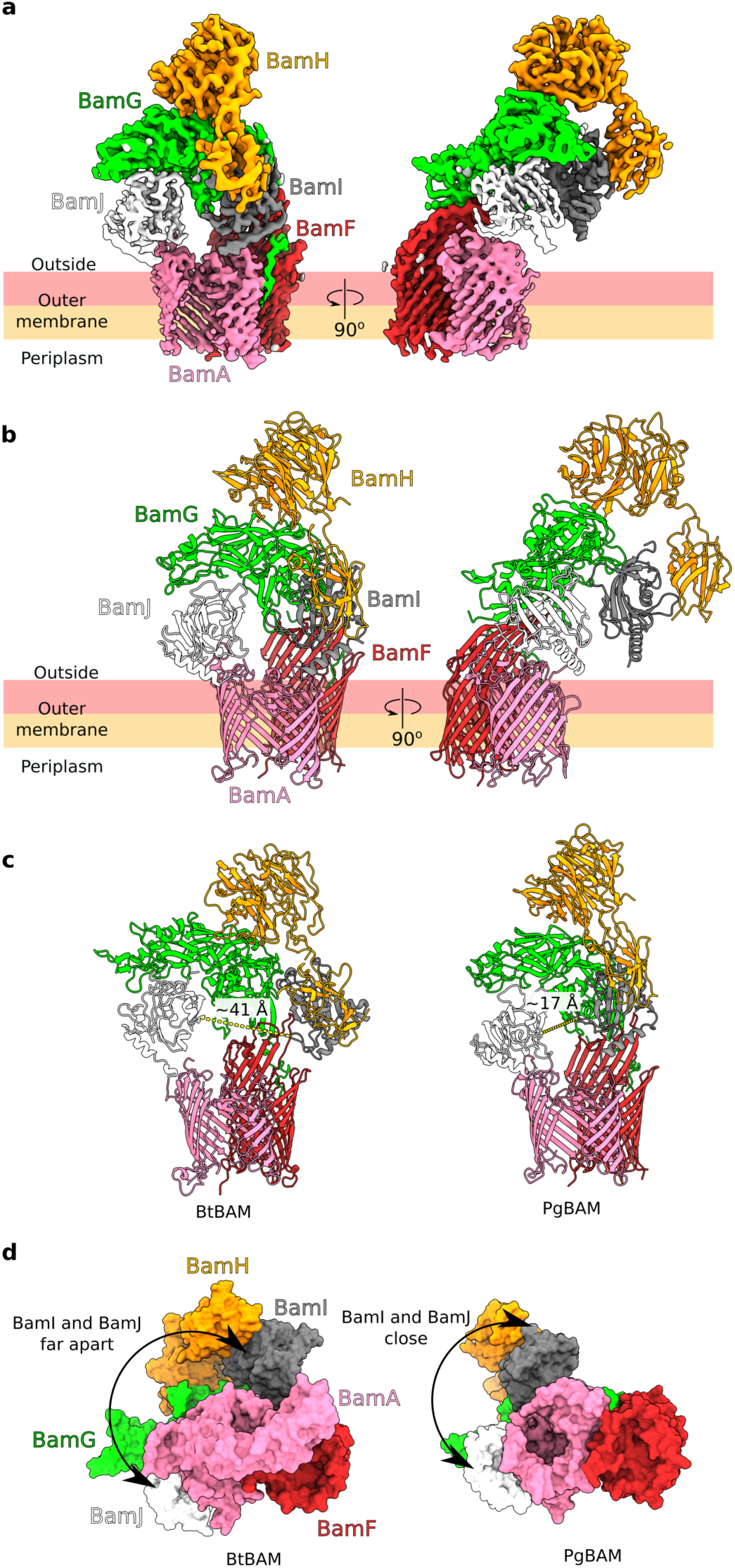
*P. gingivalis* BAM has the same architecture as BtBAM. **a**, Cryo-EM map of the PgBAM class without periplasmic density, coloured by component. **b**, PgBAM model. The colouring scheme is the same as for BtBAM. **c**, The BtBAM extracellular dome is more open than observed in the PgBAM cryo-EM structure, as demonstrated by the distance between the nearest BamI and BamJ loops (yellow dashed line). Views generated from a superposition on BamA. **d**, View of the BtBAM and PgBAM complexes from the cytoplasmic side. BtBamD is not shown for clarity. Views generated from a superposition on BamA.

The crescent-shaped BamG subunit is positioned over the BamA barrel and forms an arch-like structure approximately 60 Å above the OM surface (ED Fig. 4a). BamG interacts with BamH and BamI on one side, and with BamJ on the other. Based on its structure and the conservation of the protein core, BamG likely functions as a scaffold. The highly conserved N-terminus of BamG is wedged between two strands of the BamF barrel and interacts with the conserved BamF N-terminus, and the tri-acylated BamG lipid anchor is positioned at the BamAF interface (ED Fig. 4a,b). Together, BamF-J form a striking, half dome-shaped structure in the extracellular space, with its apex almost exactly above the lateral opening of BamA (Fig. 2d). Of the remaining three components, BamH consists of two domains. The small N-terminal domain is conserved and interacts with the BamI PPI, whereas the less-conserved C-terminal domain contacts BamG and adopts a 6-bladed β-propeller fold, with each blade consisting of 3-4 β-strands (ED Fig. 4c). Neither domain provides clues as to the potential role of BamH within the complex. The *E. coli* BAM complex also contains a β-propeller lipoprotein, BamB, facing the periplasm, but it is unclear whether the functions of the β-propellers are analogous or distinct in the two organisms. The conservation analysis of both BamI and BamJ indicates that their putative active sites are directed towards the interior of the BAM dome (ED Fig. 4d,e). A total of 17 glycosylation sites were observed in the cryo-EM density (ED Fig. 4f). Most of these are extracellular, with only one located on the periplasmic side on BamF. BamG-J all have glycosylation sites, with by far the most (10) present on BamH. All sites have the consensus D-T/S-hydrophobic motif for O-glycosylation in Bacteroidetes^37^. It is unclear if these post-translational modifications have any functional relevance.

**Figure 4.**
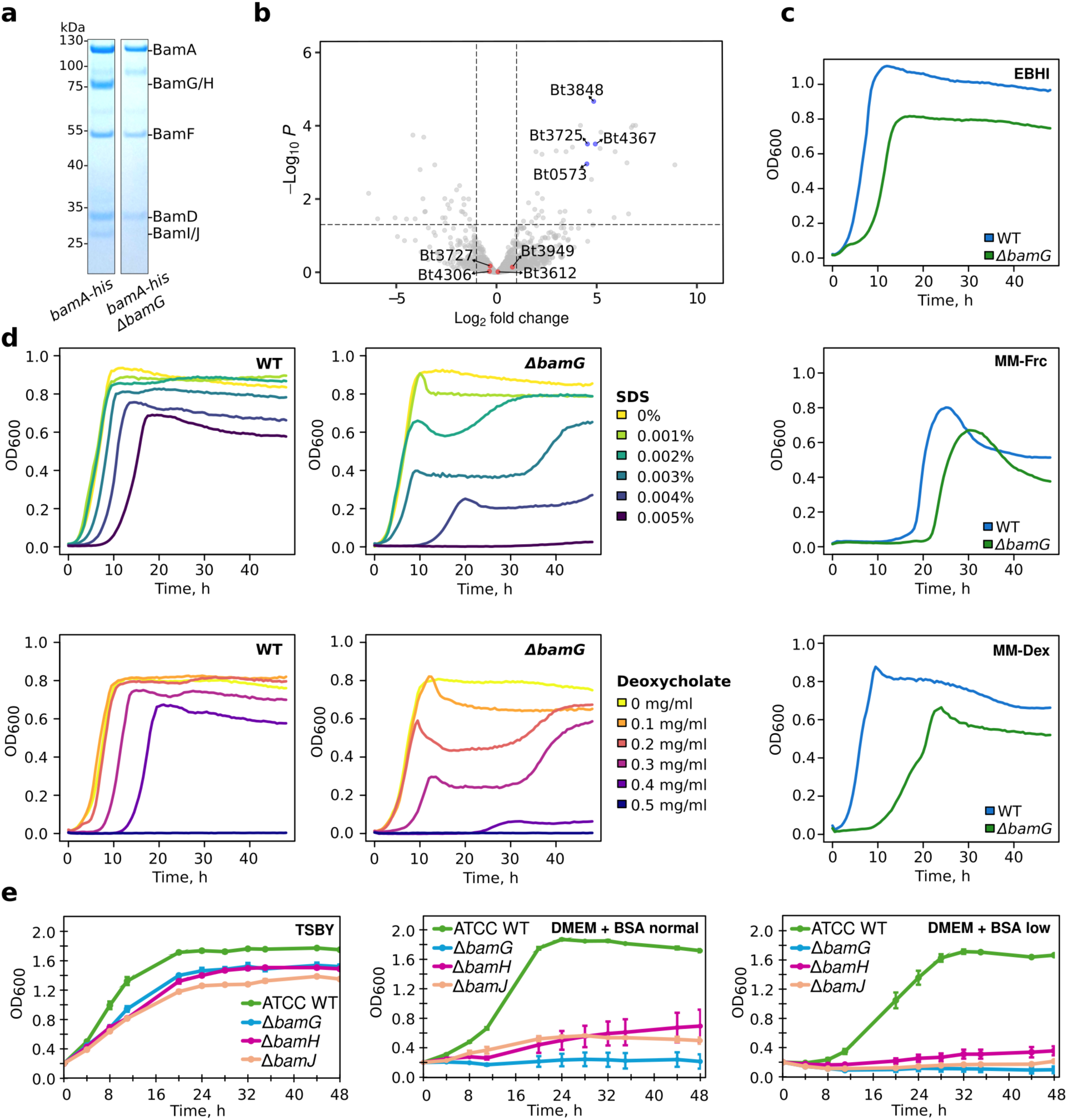
Novel BAM components are important for cell envelope integrity and growth in vitro. **a**, Coomassie-stained 4-12% SDS-PAGE gel of BtBamA_his_ pulldowns from wild-type cells and the *bamG* deletion strain. **b**, Volcano plot showing enrichment of proteins in pulldowns from *B. theta bamA_his_* 11*bamG* versus the wild type strain (n=3 biological replicates for both conditions). The vertical dashed lines represent log2 fold change=±1. Proteins above the horizontal dashed line have an adjusted p-value<0.05. **c**, Growth of wild type and 11*bamG B. theta* strains in enriched brain-heart infusion (EBHI; top), minimal medium with 0.5% fructose (MM-Frc; middle) and minimal medium with 0.4% dextran 40 and 10% enriched brain-heart infusion (MM-Dex; bottom). Each trace is an average of n=3 technical repeats. **d**, Growth of wild type and 11*bamG B. theta* strains in EBHI in the presence of indicated concentrations of sodium dodecyl sulphate (SDS; top) and sodium deoxycholate (bottom). Each trace is an average of n=3 technical repeats. **e**, Growth of *P. gingivalis* wild type, 11*bamG,* 11*bamH* and 11*bamJ* in rich medium (TSBY) and minimal medium (DMEM) supplemented with bovine serum albumin (BSA) and regular or low amounts of vitamin K (0.5 mg/l and 0.05 mg/ml, respectively) and hemin (5 mg/ml and 0.5 mg/ml, respectively). Each data point is an average of n=3 biological repeats; the error bars represent the SD.

Similarly to Proteobacterial BAM complex structures, the cryo-EM maps suggest considerable heterogeneity and/or mobility within BamA and BamD. Density for BamA POTRA domains 1-3 and for the N-terminal region of BamD is missing (residues 18-71), and the visible periplasmic components have a relatively low resolution (Supplementary Fig. 1f). Likewise, density for β1 and β2 strands of BamA is poor, suggesting heterogeneity at the lateral gate, and loose association of β1 and β16 strands (Fig. 2c and Supplementary Fig. 1e,f). The *E. coli* BamA β-barrel seam has been observed in various degrees of opening, and the seam is disordered in our BtBamA structure^1,38,39^. We also noticed increased disorder of the detergent micelle at the BamA β-barrel seam (Supplementary Fig. 2), consistent with results from molecular dynamics simulations showing membrane thinning at the *E. coli* BamA β-barrel gate^2,40^.

With regards to the BamA extracellular loops, there are notable differences between BtBamA and *E. coli* BamA (EcBamA). BtBamA has a striking, extracellular loop (EL) 3 between the β5 and β6 strands which contains 13 tyrosine residues, and which likely projects inside the dome formed by the SLPs (ED Fig. 5a-d). Sequence alignment of Bacteroidetes BamA homologues indicates that EL3 is also enriched in glycine and asparagine residues (ED Fig. 5e). There was no cryo-EM density observed for this 38-residue loop, which suggests that it samples a range of different conformations. The function of the tyrosine-rich EL3 is unclear, but its flexibility implies that it does not bind to the other BtBAM components, and it could interact with substrate β-barrels and/or, perhaps more likely given its absence in Proteobacterial BAM, SLPs. The β-barrel lumen in both BtBamA and EcBamA is occluded mainly by EL6, but these loops differ in length and structure, with BtBamA EL6 being much shorter (ED Fig. 5a-c). Both proteins have a short, partially α-helical EL corresponding to EL5 in BtBamA and EL4 in EcBamA. BtBamA EL5 contacts the BamJ PPI, but the functional significance of this is unclear.

**Figure 5.**
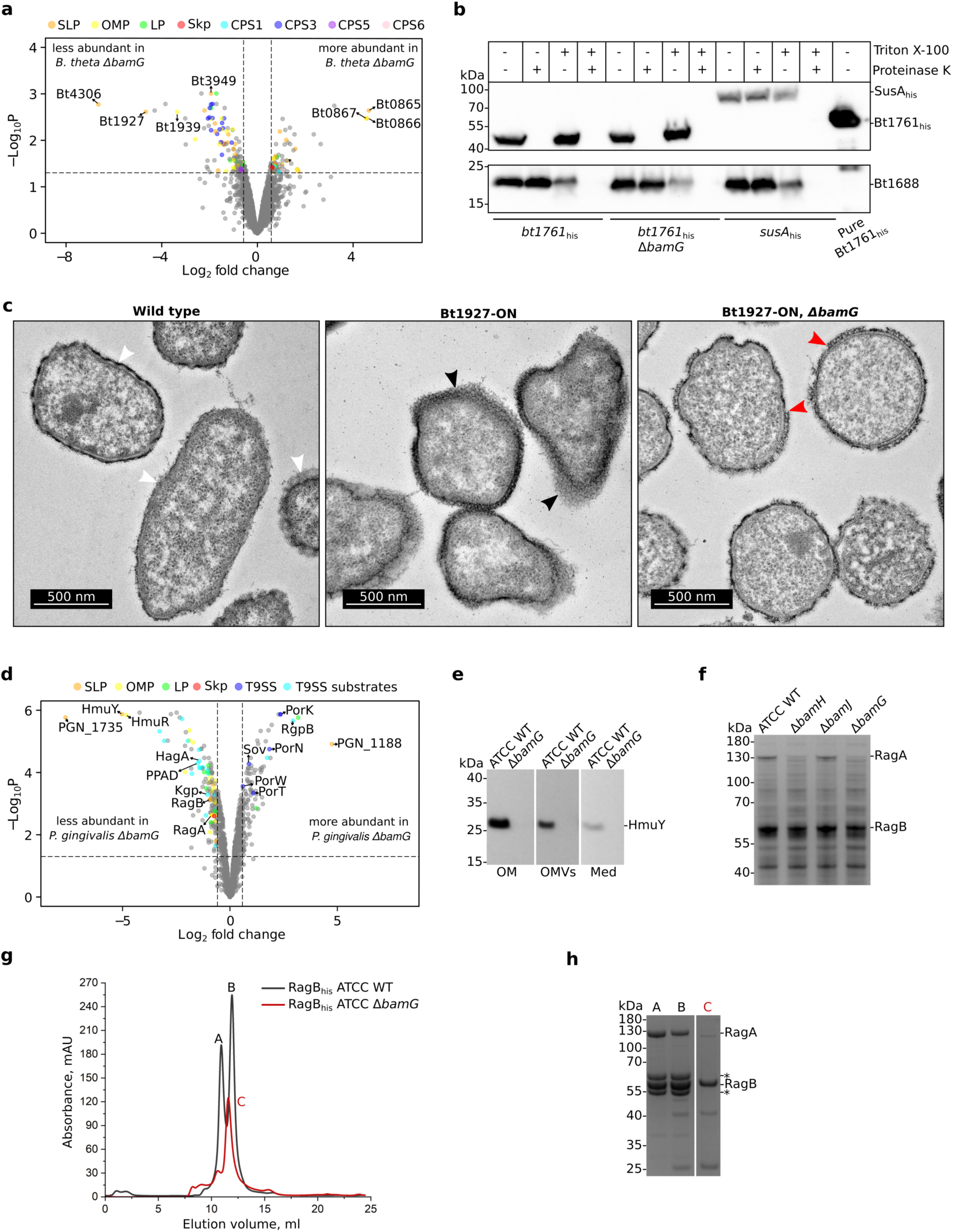
Loss of extracellular BAM components results in pleiotropic changes at the cell surface. **a**, Volcano plot showing wild type *B. theta* vs 11*bamG* quantitative proteomics results (n=3 biological replicates for each). The vertical dashed lines represent log2 fold change = ±0.58. Proteins above the horizontal dashed lines have an adjusted p-value < 0.05. Data points of interest are labelled and colour-coded according to function or cellular localisation. **b**, *B. theta* proteinase K shaving assay for testing protein surface exposure. The top immunoblot was probed with anti-His-HRP conjugate. The bottom panel shows the same immunoblot re-probed with StrepTactin-HRP. Bt1688 is a cytoplasmic biotin-binding protein. SusA is a periplasmic neopullulanase which was used as an OM lysis control. **c**, Transmission electron microscopy images of thin sections of fixed *B. theta* cells at 13,500x magnification. The white arrowheads point to the capsule which varies in thickness between cells. The black arrowheads point to the S-layer in Bt1927-ON cells. The red arrowheads point to the discontinuous S-layer in the Bt1927-ON, Δ*bamG* strain. **d**, Volcano plot showing wild type *P. gingivalis* vs Δ*bamG* quantitative proteomics results (n=3 biological replicates for each). **e**, Anti-HmuY immunoblot of outer membrane (OM), OM vesicle (OMV) and culture medium (Med) fractions of wild type and 11*bamG P. gingivalis* strains. **f**, SDS-PAGE analysis of *P. gingivalis* wild type, 11*bamH,* 11*bamG,* and Δ*bamI* strain whole cell lysates. **g**, RagB_his_ pulldowns from *P. gingivalis* wild type and 11*bamG* strains analysed by size exclusion chromatography on a Superdex 200 Increase 10/300 GL column. **h**, SDS-PAGE analysis of indicated peaks from (**g**). The asterisks denote bands from peaks A and B that correspond to singly-cleaved RagA, confirmed by mass spectrometry. SLP, surface-exposed lipoprotein; OMP, outer membrane protein; LP, lipoprotein; CPS, capsular polysaccharide; T9SS, type 9 secretion system; PPAD, peptidylarginine deiminase.

### The BtBAM architecture is conserved in *Porphyromonas gingivalis*

To obtain more information on the structural conservation of the BAM complex we turned to *Porphyromonas gingivalis* strain ATCC 33277, a prominent, well-studied oral pathogen belonging to the family Porphyromonadaceae. Compared to the large genomes of *Bacteroides* spp. (*B. theta*; ∼6.2 Mb, ∼4800 ORFs), *P. gingivalis* has a much smaller genome (∼2100 ORFs), making it an interesting organism to assess the conservation of the BAM complex. Tagging PgBamA at the N-terminus led to low yields of intact complex due to proteolytic degradation of the periplasmic domain, presumably by the ubiquitous gingipain surface proteases characteristic of *P. gingivalis*^41^. A tag added to the C-terminus of PgBamG (PGN_1735) allowed us to purify the intact complex and determine its cryo-EM structure at ∼3.2 Å resolution (Fig. 3a,b, Supplementary Fig. 3 and Supplementary Table 1). The composition and architecture of the PgBAM complex is very similar to that of *B. theta*, with a pronounced extracellular dome formed by BamF-J. Only very weak, not modelled, density was observed for the periplasmic components, possibly due to clipping by gingipains, similar to what was observed in the isolation of tagged PgBamA (ED Fig. 6a). All components of PgBAM and BtBAM are similar in size except for BamG, which is 88 residues longer in *B. theta*. Structural alignments of individual BtBAM and PgBAM components show high structural similarity for BamAFGIJ (ED Fig. 7). The highest Cα-Cα RMSD of 6.3 Å was observed for BamH, though both PgBamH and BtBamH consist of an N-terminal Ig-like domain and a C-terminal 6-bladed β-propeller. The extracellular dome of PgBAM is much more compact than that of BtBAM, as indicated by the distance between the BamI and BamJ PPIs (Fig. 3c,d and Supplementary Movie 1). A total of 12 glycosylation sites are present within PgBAM, mostly located on BamG (7 sites) (ED Fig. 6b). The tri-acylated PgBamG lipid anchor is clearly defined and, like in *B. theta*, positioned at the BamAF interface (ED Fig. 6c). EL3 in *P. gingivalis* BamA is 39 residues long and contains 12 tyrosines. Similar to BtBamA, there is no cryo-EM density for this loop in PgBamA.

### BAM surface lipoproteins are important for growth and OM integrity

Next, we constructed a *B. theta bamG* deletion strain to investigate its function. Given its likely role as a scaffold protein, we hypothesised that BamG removal might result in loss of all SLP components of the BAM complex (BamG-J). Indeed, BtBamA_His_ pulldowns from a τι*bamG* background confirmed this notion (Fig. 4a). The τι*bamG* strain showed growth defects in rich medium and minimal medium with both monosaccharides and polysaccharides as carbon sources (Fig. 4c). The deletion of *bamH* resulted in an intermediate phenotype (Supplementary Fig. 4), suggesting that the defect observed in the τι*bamG* strain is due to a cumulative loss of BamH, BamI, BamJ and possibly BamG itself. When *B. theta* was cultured in the presence of the OM stressors sodium dodecyl sulphate (SDS) and sodium deoxycholate, we observed a severe decrease in OM stress tolerance in the τι*bamG* strain compared to the wild-type strain (Fig. 4c). Thus, our data show that BamG is important *in vitro*, but not essential. Alternatively, its paralogue Bt1786 might be able to partially complement the lack of BtBamG. Bt1786 is in a putative operon with Bt1785, a BtBamF paralogue, and the Bt1785-6 complex model predicted by AF2 is similar to the BamFG unit in our BtBAM cryo-EM structure (Supplementary Fig. 5). However, we have not been able to detect Bt1785 or Bt1786 in our proteomics data, including those for τι*bamG* strains, arguing against the existence of an alternative BtBAM complex involving Bt1785-86 that could rescue phenotypes in the τι*bamG* strain.

For *P. gingivalis*, growth of *bamG*, *bamH* and *bamJ* deletion strains in rich medium was only slightly reduced (Fig. 4e). However, we observed a drastic growth reduction in minimal medium (DMEM containing 1% bovine serum albumin as carbon and nitrogen source) when supplemented with standard concentrations of vitamin K (0.5 mg/L) and hemin (5 mg/L) (Fig. 4e; DMEM + BSA normal). Growth was further compromised when the concentration of supplements was decreased 10-fold (Fig. 4e; DMEM + BSA low). Notably, the *bamH* and *bamJ* mutants remained viable after 48 hours, whereas the *bamG* mutant was dead shortly after inoculation. Moreover, the growth defect in minimal medium for PgBAM mutants was much more pronounced than for mutants of the RagAB OM β-barrel-SLP complex involved in peptide uptake^42^. This suggested that the effect of these mutations on growth might result from a cumulative impairment of SLP-barrel complex assembly in the OM.

### Deletion of *bamG* affects cell surface structure biogenesis in *B. theta*

We reasoned that deletion of *bamG* may have resulted in changes to the proteome that were responsible for the OM integrity defects. To compare the relative abundance of membrane proteins in *B. theta* wild type and the τι*bamG* strain, we employed quantitative proteomics using intensity-based absolute quantification (iBAQ)^43^ on total membrane fractions. A total of 131 proteins were less abundant (log_2_ fold change<-0.58; p<0.05) in the τι*bamG* strain compared to wild type *B. theta* (Fig. 5a and Supplementary Table 3). Pulldown experiments showed that deletion of *bamG* leads to loss of all SLPs from the BtBAM complex (Fig. 4a,b), but the proteomics show that only the BamG (Bt4306) and BamJ (Bt3949) SLP levels are lower in the τι*bamG* strain (Fig. 5a), while levels of other BtBAM components are not affected (Supplementary Table 3). In addition, most proteins from the Bt0598-Bt0614 capsular polysaccharide (CPS) locus 3^44^ show lower levels in the τι*bamG* strain. CPS3 is the main capsule produced during *in vitro* growth of *B. theta*^45^, and defects might explain the increased sensitivity of the τι*bamG* strain to OM stress (Fig. 4d). In total, 14 OM β-barrels had lower abundance in the τι*bamG* strain, nine of which are TBDTs; 35 lipoproteins were downregulated in the τι*bamG* strain, 24 of which are likely surface exposed based on the presence of a LES. On the other hand, the 76 proteins that were enriched (log_2_ fold change>0.58; p<0.05) in *B. theta* τι*bamG* include 8 SLPs and 4 OM β-barrels (3 TBDTs). The Bt0865 SusC-like TBDT of unknown function, Bt0866 SusD-like SLP and Bt0867 SLP were the most highly enriched proteins in the total membrane fraction, which suggests that both OM β-barrel and SLP biogenesis are not inhibited by lack of the BtBAM SLP components. The periplasmic OMP chaperone Skp (Bt3724) was also slightly enriched in the *bamG* deletion strain.

The proteins with altered abundance in the *bamG* deletion strain relative to wild type represent relatively small fractions of the total number of β-barrels and (S)LPs. The proteomics data therefore show that the BtBAM complex missing all its SLP components does not result in global depletion of OM β-barrels and SLPs, and it does not drastically affect the normal BtBAM β-barrel insertase function *in vitro*. However, the proteomics data do not provide information on whether SLPs reach the cell surface in the τι*bamG* strain. To test this, we used a proteinase K shaving assay on the abundant surface glycan binding lipoprotein Bt1761^46^ in wild-type and τι*bamG* cells, and we found that Bt1761 is still surface-exposed in the deletion strain (Fig. 5b). This suggests that the extracellular BtBAM dome is not required for SLP flipping.

The Bt1926-27 locus, which encodes a 100 kDa phase-variable SLP S-layer protein (Bt1927)^47,48^,is much less abundant in the *B. theta* τι*bamG* strain. Its putative OM anchor or secretion partner (Bt1926)^47^, predicted to form an 8-stranded β-barrel, was not detected in the τι*bamG* strain and is thus absent from the volcano plot in Fig. 5a. Transmission electron microscopy images of thin sections of wild-type *B. theta* show that the cells are surrounded by a diffuse density of variable width, which likely corresponds to the capsule, but there is no density attributable to an S-layer (Fig. 5c). This is because most wild type *B. theta* cells do not transcribe *bt1927* as the upstream invertible promoter is inactive, but the promoter can be engineered to make all cells produce the S-layer^47^. Micrographs of the engineered strain producing Bt1927 (Bt1927-ON) clearly show cells surrounded by an electron-dense, lattice-forming structure that corresponds to the S-layer (Fig. 5c and ref. 47). The S-layer appears to be loosely associated or detaching from some cells, but it is unclear if this is real or an artefact of sample preparation. When S-layer production is turned on in a *bamG* deletion background, *B. theta* only forms a partial, discontinuous S-layer, observed as patchy, electron-dense structures in micrographs (Fig. 5c). Thus, the extracellular dome of BtBAM clearly plays a role in the biogenesis of at least some cell surface structures, potentially by acting as an assemblase that promotes association of Bt1927 with its putative anchor or secretion partner Bt1926^47^.

### Deletion of *bamG* disrupts the main SLP-barrel complexes of *P. gingivalis*

Since the genome of *P. gingivalis* encodes a relatively low number of proteins, we used whole cells instead of membrane fractions for quantitative proteomics. 1,500 proteins were identified, providing excellent proteome coverage of 71%. The level of 102 proteins was significantly increased (log_2_ fold change > 0.58) whereas the level of 146 was significantly decreased (log_2_ fold change < −0.58) (Fig. 5d and Supplementary Table 4). BamG (PGN_1735), as expected, was the most depleted protein. The levels of other components of PgBAM remained unchanged, except for BamJ (PGN_1188), which intriguingly was the most enriched protein in the analysis. This suggests that the levels of the PgBAM SLPs are unaffected in the *bamG* mutant, even though they are likely not associated with BamADF.

There are two experimentally confirmed SLP-barrel complexes in *P. gingivalis*: the peptide transporter RagAB^42^ and the heme uptake system HmuYR^49^. The levels of both the HmuY SLP and the HmuR TBDT are severely decreased in the τι*bamG* strain (Fig. 5d and Supplementary Table 4). To verify the proteomics data, we probed western blots with anti-HmuY SLP antibodies and observed that HmuY was undetectable in the OM, outer membrane vesicles (OMVs) and culture medium in the τι*bamG* strain (Fig. 5e). The levels of the RagA OM β-barrel and the RagB SLP were moderately decreased to a similar extent in proteomics, even though RagA was barely visible on SDS-PAGE of a τι*bamG* lysate compared to RagB (Fig. 5f). To check if the RagAB complex is assembled in the τι*bamG* strain, we isolated His-tagged RagB from a τι*bamG* background (Fig. 5g). The SDS-PAGE profile revealed a clear RagB band, but almost no full-length RagA (Fig. 5h). This suggests that RagA undergoes proteolysis due to the impaired assembly of the RagAB complex. This is consistent with a previous observation that the level of RagA is also much lower in a τι*ragB* mutant^42^. Among the proteins with significantly reduced levels in the τι*bamG* strain are 17 lipoproteins, 9 TonB-dependent transporters (TBDTs), and 9 other β-barrels (Fig. 5d and Supplementary Table 4).

Notably, the deletion of *bamG* also has a pronounced effect on the T9SS. The levels of T9SS OM core components Sov, PorN, PorK, PorT, and PorW are significantly increased, although PorV remains unchanged (Fig. 5d and Supplementary Table 4). Additionally, the periplasmic T9SS components PorL and PorM, are also unchanged. Despite increased levels of core T9SS components, the levels of 19 out of 35 T9SS substrates (including RgpA and Kgp gingipains, peptidylarginine deiminase, and hemagglutinin A) are significantly lower in the *bamG* deletion strain (Fig. 5d and Supplementary Table 4). In addition, there is one 8-stranded beta barrel (PGN_0297, significantly increased) in the experimentally confirmed BamA operon (PGN_0296-PGN_0301)^50^, which is potentially involved in the T9SS. A deletion mutant of PGN_0297 is completely white on blood agar plates, like T9SS mutants. Intriguingly, PGN_0297 is the neighbour of BamH (PGN_0296; BamA corresponds to PGN_0299). Clearly, the multidirectional effects of *bamG* deletion on the T9SS warrant further investigation. Overall, out of 146 proteins with significantly reduced levels, there are 8 inherently essential proteins and 66 essential proteins for fitness in both hTERT-immortalized human gingival keratinocyte (TIGK) cell model and murine abscess model^51,52^ (Supplementary Table 4). Cumulatively, about half of the proteins with decreased level in the *bamG* mutant are important for pathophysiology of *P. gingivalis*^51,52^.

## Discussion

To date, only a very limited set of non-canonical BAM-associated lipoproteins has been identified^53^, with unknown functions and associated with BamA in the periplasmic space. Here, we have discovered a completely novel architecture of the BAM machinery in the Bacteroidetes phylum (ED Fig. 8a). Besides the minimal core formed by BamA and BamD, it also includes the essential β-barrel BamF, which serves as an attachment platform for an extracellular dome-like structure composed of four SLPs, BamG, BamH, BamI, and BamJ. Given that the defining difference between OMPs of Bacteroidetes and Proteobacteria is the dominance of SLPs in the former, we propose that the novel components and architecture of Bacteroidetal BAM have evolved to enable the efficient biogenesis of SLPs and SLP-barrel complexes (ED Fig. 8b).

How the ubiquitous SLPs of Bacteroidetes reach the cell surface is an important, unresolved question. In Proteobacteria, there are several systems for which a function in secretion and surface exposure of lipoproteins has been proposed, such as the surface lipoprotein assembly modulator (Slam)^54^, the type 2 secretion system in *Klebsiella* spp.^55^, the Aat system for secretion of unique, glycine-acylated lipoproteins^56^, and the OM LPS assembly protein LptD in *B. burgdorferi*^57^. With the exception of LptD, *B. theta* and some other Bacteroidetes lack all of the aforementioned systems while its genome encodes > 400 SLPs. Given the central roles of SLPs in Bacteroidetes OM biology, it is very likely that the SLP flippase is essential. However, our BamA pulldowns did not identify any OMP that satisfied the criteria of essentiality and having a predicted structure compatible with a protein flippase function (*i.e*., a barrel with a lateral opening). Moreover, no essential OMP besides BamA (Bt3725) itself has been reported that satisfies those criteria^58–60^. We therefore propose that BamA is the Bacteroidetes SLP flippase. There is precedent from Proteobacteria for such a role, since the stress sensor lipoprotein RcsF in *E. coli* is exported to the cell surface by EcBamA. In a so-called push-and-pull mechanism, RcsF interacts with EcBAM on the periplasmic side and its secretion and surface exposure are triggered by incoming OMP substrates^61,62^. It is an intriguing possibility that Bacteroidetes SLPs might use hybrid β-barrels formed by BamA and substrate OMPs as a secretion pathway for surface exposure. While it is unclear whether SLPs are secreted (partially) folded or unfolded, hybrid TBDT-BamA barrels might be large enough to allow passage of folded SLPs or SLP domains.

BtBamA and PgBamA are structurally very similar to EcBamA, including the 5 POTRA domains, the EL6 cap, and the dynamic β-barrel seam between strands 1 and 16. Furthermore, Bacteroidetes β-barrel proteins have an aromatic β-signal at their C-terminus. This suggests that BamA-mediated substrate barrel folding and OM insertion in Bacteroidetes follows the same “hybrid-barrel budding” mechanism as that described for *E. coli*. The most pronounced difference between Bacteroidetal BamA and Proteobacterial BamA is the tyrosine-rich EL3. Recently, it has been shown that bacteroidetocins, antibacterial peptide toxins produced by Bacteroidetes, target BamA EL3^63^, which suggests that this loop is critical to BAM function. The YG-rich composition of EL3 is somewhat reminiscent of nuclear pore and peroxisomal import proteins, where FG- and YG-rich domains are proposed to form a hydrogel mesh via which cargo is imported^64,65^. We speculate that the BtBamA EL3 might interact with lipoproteins directed to the cell surface to facilitate their flipping and OM insertion.

The role of the integral OMP BamF in the Bacteroidetes BAM complex is very intriguing. Given BamF appears to be essential whereas simultaneous loss of all SLPs in the *bamG* deletion is tolerated *in vitro*, it is unlikely that the sole function of BamF would be to provide, via BamG binding, a physical link between BamAD and the extracellular dome formed by BamG-J. It is possible that BamF, via its extensive interface with BamA, is required for the stability of BamA analogous to the role of LptE within the LPS insertase LptDE^66^. Alternatively, or perhaps in addition, BamF could contribute functionality to the BamAD core insertion complex via, *e.g*., providing a positively charged binding site for the acidic LES of SLPs (ED Fig. 9) or via interactions with extra-membrane regions of BamA.

The structure and positioning of the extracellular dome formed by BamG-J, and the involvement of two PPIs (BamIJ), suggests that this structure may form an extracellular chaperone or assembly cage for the formation of SLP-barrel complexes that are a hallmark of the Bacteroidetes phylum. This hypothesis is supported by the fact that the cage is large enough to accommodate folded SusCD complexes (ED Fig. 10). The removal of this cage in τι*bamG* led to substantial growth and OM defects and changes in OMP abundance in both *B. theta* and *P. gingivalis*. Loss of BamG in *B. theta* resulted in reduced expression of capsule biosynthesis genes, potentially explaining why the τι*bamG* strain is much more sensitive to OM stressors. *B. theta* lacking BamG is also incapable of producing a complete S-layer, possibly due to impaired attachment of the S-layer SLP Bt1927 to its putative anchor OM β-barrel Bt1926. Loss of BamG in *P. gingivalis* led to low levels of heme and peptide uptake systems, which explains the dramatic growth defects observed in minimal medium with bovine serum albumin used as the carbon source. In addition, *P. gingivalis* τι*bamG* showed decreased levels of T9SS substrates even though many components of the T9SS machinery were more abundant, suggesting that some aspect of secretion is affected in the mutant strain. While the observed proteome changes may seem modest both for *B. theta* and *P. gingivalis*, it should be noted that they result from removal of extracellular BAM components only. Moreover, the proteomics report on total protein abundance and do not provide clues about, *e.g*., defective or slower assembly of SLP-barrel complexes. Finally, it has been shown that BamG is one of only ∼80 core fitness components for mouse gut colonisation by several *Bacteroides* spp.^58^, suggesting that the effects of BamG removal would be more dramatic *in vivo*. Together, our functional data suggest that the extracellular dome plays an important role in OMP complex and cell surface structure biogenesis in Bacteroidetes through an unknown mechanism.

Given its essential role in OMP biogenesis and maintaining bacterial cell integrity, BAM has emerged as an important target for antibacterial therapy. Currently, several types of BAM inhibitors have been developed, including small molecules, peptides, natural products, and antibodies^67^. The mode of action of these compounds involves binding to critical regions of BamA (such as L3, L4, L6 loops and lateral seam) or interfering with the BamA-BamD interface, thereby impairing proper assembly of the BAM complex and folding of OMPs^63,68–70^. To reach the periplasmic components of BAM, potential inhibitors must be able to cross the OM, which drastically limits the number of useful compounds^71^. Bacteroidetes BAM is a promising target for inhibitors due to the extracellular localization of the BamG-J dome, which makes it an easily accessible target. Moderate conservation of these surface-exposed components provides an opportunity to discover inhibitors targeting pathogenic Bacteroidetes such as *P. gingivalis* while avoiding the commensals.

In conclusion, our work provides a foundation to formulate hypotheses to investigate and establish the function of the Bacteroidetes BAM complex, and to dissect the roles of the individual components. Analogous to the work on Proteobacterial BAM over the past two decades, we expect this to yield exciting insights that will provide a more comprehensive understanding of outer membrane biogenesis in Gram-negative bacteria.

## Methods

### Bacterial strain culturing and genetic manipulation

Bacterial strains used in this study are listed in Supplementary Table 5. *E. coli* strains were grown at 37 °C in lysogeny broth (LB, 20 g/l) with 100 µg ml^−1^ ampicillin and on 1.5% agar plates. For constructing *B. theta* chromosomal mutants and protein production, strains were cultured in enriched brain-heart infusion (EBHI), containing 37 g/l Oxoid brain-heart infusion powder, 5 g/l yeast extract and 1 μg/ml hemin, supplemented with 0.5 g/l cysteine and 1 μg/ml vitamin K. For growth curve and proteinase K experiments, defined minimal medium (MM) supplemented with 1 μg/ml hemin and 0.5% carbon source was used, unless otherwise indicated. Bacteria were grown under anaerobic conditions at 37°C in a Don Whitley A35 workstation. *B. theta* genetic deletions and mutations were created by allelic exchange using the pExchange-tdk vector^72^. Briefly, the constructed pExchange-tdk plasmids, containing the mutations/deletions plus ∼700 bp flanks up- and downstream (DNA inserts were synthesised by Twist Bioscience), were transformed into S17 λ pir *E. coli* cells, to achieve conjugation with the *B. theta* recipient strain. The conjugation plates were scraped and *B. theta* cells that underwent a single recombination event were selected for by plating on BHI-hemin agar plates containing gentamicin (200 μg/ml) and erythromycin (25 μg/ml). Eight colonies were restreaked on fresh BHI-hemin-gent-ery plates. Single colonies were cultured in BHI-hemin overnight and pooled. To select for the second recombination event, pooled cultures were plated on BHI-hematin agar plates containing 5-fluorodeoxyuridine (FUdR; 200 μg/ml). After 48-72h of growth, FUdR-resistant colonies were restreaked on fresh BHI-hematin-FUdR. From these, single colonies were cultured in BHI and genomic DNA was extracted and screened for the correct mutations using diagnostic PCR. This procedure was used to introduce the N-terminal His_7_-tag on BtBamA, and *bamG* and *bamH* deletions into the chromosome of *B. theta* VPI-5482 *tdk^−^* and BamA_his_, as well as a *bamG* deletion in the Bt1927-ON background.

*P. gingivalis* strains were grown anaerobically (90% nitrogen, 5% carbon dioxide and 5% hydrogen) at 37°C in enriched tryptic soy broth (eTSB; 30 g/l tryptic soy broth, 5 g/l yeast extract, 5 mg/l hemin, 0.5 mg/l menadione and 0.25 g/l L-cysteine) or on eTSB blood agar plates with 1.5% agar and 5% defibrinated sheep blood. For mutant selection, appropriate antibiotics were added: erythromycin at 5 μg/ml or tetracycline at 1 μg/ml. *P. gingivalis* mutants were generated through homologous recombination. For deletion mutants, 1 kb fragments flanking the *bamG*, *bamH* or *bamJ* genes and chosen antibiotic resistance cassettes were amplified by PCR and cloned into pUC19 plasmid using restriction cloning. For expression of C-terminally-His-tagged BamG protein, master plasmid was first generated. 1 kb fragment coding the C-terminal part of BamG and 1 kb fragment downstream of it as well as an antibiotic resistance cassette were amplified and cloned into pUC19 plasmid using restriction cloning. The 7His-tag was then introduced into the master plasmid by PCR. Plasmids used for mutant construction are listed in Supplementary Table 6. Plasmids were electroporated into *P. gingivalis* competent cells, followed by plating the cells on eTSB blood agar with appropriate antibiotics and growth for 10 days. All generated plasmids were confirmed by PCR and Sanger sequencing. *B. theta* and *P. gingivalis* clones were screened by PCR and verified by Sanger sequencing.

### BtBAM expression and purification

The *B. theta bamA_his_* strain was cultured overnight in EBHI and used to inoculate 10 litres of rich medium (25 g/l brain-heart infusion powder (Oxoid); 3.5 g/l yeast extract; 1 µg/ml vitamin K; 0.5 g/l L-cysteine) that was equilibrated at 37°C overnight in an anaerobic chamber. 3 ml of overnight culture was used per 500 ml bottle of pre-warmed medium. The cultures were grown for 5-7 hours until late exponential phase (OD_600_∼1.8-2.0). Cultures were pelleted by centrifugation at 6,000g for 30 min at 4°C, and the pellets were stored at −20°C.

Cell pellets were processed in 2 litre batches to maximize yield. Cells were thawed, homogenised in Tris-buffered saline (TBS; 20 mM Tris-HCl pH 8.0, 300 mM NaCl), supplemented with DNase I, and lysed by a single pass through a cell disruptor at 22-23 kpsi. The total membrane fraction was isolated by ultracentrifugation at 200,000g for 50 min at 4°C. Membranes were homogenised and solubilised in 30 ml TBS with 0.75% dodecyl maltoside (DDM) and 0.75% decyl maltoside (DM) for 1 h at 4°C with stirring. Insoluble material was pelleted by ultracentrifugation at 200,000g for 30 min at 4°C. The supernatant was loaded on a 1.5 ml chelating Sepharose gravity flow column charged with Ni^2+^ ions at 4°C and equilibrated with TBS supplemented with 0.15% dodecyl maltoside (DDM). The column was washed with 30 column volumes (CV) TBS supplemented with 30 mM imidazole and 0.15% DDM, and bound material was eluted with 4 CV TBS supplemented with 200 mM imidazole and 0.15% DDM. The eluate was concentrated using an Amicon Ultra filtration device (100 kDa MWCO) and loaded on a Superdex 200 Increase 10/300 GL column equilibrated in 10 mM HEPES-NaOH, 100 mM NaCl, 0.03% DDM pH 7.5. Peak fractions were analysed by SDS-PAGE and Coomassie staining for purity. Gel bands of interest were cut out and identified by peptide fingerprinting at the Metabolomics and Proteomics Laboratory, University of York. Material purified from five separate 2 litre batches was combined and subjected to a final size exclusion chromatography run as above. The BtBAM complex was concentrated, flash-frozen in liquid nitrogen, and stored at −80°C.

### BtBAM cryo-EM structure determination

Purified BtBAM complexes at 8 mg/ml were applied to glow-discharged Quantifoil R1.3/1.2 holey carbon 200 mesh Cu grids, followed by blotting and plunge-freezing in liquid ethane using a Vitrobot Mark IV device at 4°C and 100% humidity. Data were collected on a Titan Krios electron microscope operating at 300 kV accelerating voltage on a Falcon 4i direct electron detector with a Selectris energy filter (10 eV slit). Movies were recorded in EER format at a magnification of 165,000, corresponding to a sampling rate of 0.74 Å/pixel. The defocus was varied between −0.8 µm and −2.0 µm during data collection. The total dose for each movie was ∼35 e^−^/Å^2^. 13,558 movies were collected in total. Data collection parameters are summarised in Supplementary Table 1.

The cryo-EM data processing workflow is shown in Supplementary Fig. 1. All data processing was done in cryoSPARC v4.5.3^73^. Movies were patch-motion-corrected, followed by patch CTF correction. Particles were picked manually to generate 2D classes for template-based picking. 2,266,553 particles were extracted in 224×224 pixel boxes (1.48 Å/pixel) and subjected to two rounds of 2D classification. Particles from 2D classes resembling protein density were pooled into a single particle stack containing 308,834 images, which was then used for ab initio reconstruction with 4 classes. 2D classes containing 57,300 noise and junk particles were used for ab initio reconstruction of 4 decoy classes. The 308,834-particle stack was subjected to heterogeneous refinement using the 8 ab initio classes as initial volumes. 137,389 particles from the best two classes from the first round of heterogeneous refinement were subjected to another round of heterogeneous refinement against the same initial volumes as in the first round. This resulted in two classes with clear protein features: class 1 with 49,137 particles with no periplasmic density and class 2 with 55,224 particles and periplasmic density. Particles belonging to these two classes were independently used in non-uniform refinement^74^, yielding 3.41 Å (class 1) and 3.49 Å (class 2) reconstructions. Refined particles were re-extracted in 448×448 pixel boxes (0.74 Å/pixel) and subjected to another round of non-uniform refinement with enabled per-particle defocus and global CTF parameter (tilt and trefoil) refinement. 48,843 particles from class 1 yielded a final reconstruction at 3.28 Å; 56,321 particles from class 2 yielded a final reconstruction at 3.46 Å. Global resolution was estimated using gold-standard Fourier shell correlation curves and the 0.143 criterion (Supplementary Fig. 1c). Particle viewing direction distribution plots and local resolution estimates are displayed in Supplementary Fig. 1d-f.

BtBamGHIJ AF2 models were docked into the class 1 volume and subjected to cycles of manual adjustment in Coot^75^ and real space refinement in Phenix^76^. Mannose monosaccharides were placed in O-glycan densities to prevent the protein model from moving into the glycan density. Similarly, BtBamADF AF2 models were docked, manually adjusted and refined against the class 2 volume. The refined BtBamGHIJ class 1 model was rigid-body-refined against the class 2 volume, which had much poorer density for the BtBamGHIJ region than class 1 (Supplementary Fig. 1f), to obtain the complete BtBAM model. A composite map incorporating the best parts of the volume representing each class was generated using phenix.combine_focused_maps. Model refinement statistics are shown in Supplementary Table 1.

### PgBAM cryo-EM structure determination

Purified PgBAM complexes at 4.9 mg/ml were applied to glow-discharged Quantifoil R2/1 holey carbon Cu grids (200 mesh), followed by blotting and plunge-freezing in liquid ethane using a Vitrobot Mark IV (Thermo Fisher) set to 95% humidity and 4°C with the following blotting parameters: 2 s blot time, 0 s wait time, blot force −4, 0 s drain time and a total blot of 1. Data were collected using a Titan Krios G3i (Thermo Fisher; Solaris, Poland) electron microscope operating at 300 kV accelerating voltage on a Gatan K3 Summit direct electron detector with a Gatan Quantum energy filter. Movies were recorded in TIFF format at a magnification of 105,000x, corresponding to a sampling rate of 0.84 Å/pixel. The defocus was varied between −0.6 μm and −1.5 μm during data collection. The total dose for each movie was ∼41 e^−^/Å^2^. 11,459 movies were collected in total. Data collection parameters are summarised in Supplementary Table 1.

The cryo-EM data processing workflow is shown in Supplementary Fig. 3. All data processing was done in cryoSPARC v4.5.3. Movies were patch-motion-corrected, followed by patch CTF correction. 3 rounds of 2D classification were performed to generate 2D classes for template-based picking. 4,830,281 particles were extracted in 200×200 pixel boxes (1.68 Å/pixel) and subjected to two rounds of 2D classification. Particles from 2D classes resembling protein density were pooled into a single particle stack containing 349,717 images, which was then used for ab initio reconstruction with 4 classes. 2D classes containing 9,273 noise and junk particles were used for ab initio reconstruction of 3 decoy classes. Particles were re-extracted in 384×384 pixel boxes (0.84 Å/pixel). The 104,912-particle stack was subjected to heterogeneous refinement using the 5 ab initio classes as initial volumes. 101,685 particles from the best two classes were combined and subjected to ab initio reconstruction (2 classes) with class similarity parameter set to 0.0001. The class containing 86,584 particles was used in non-uniform refinement, yielding a 3.45 Å reconstruction. Refined particles were subjected to Local CTF refinement and Global CTF refinement and used again in non-uniform refinement, yielding a final reconstruction at 3.24 Å (PgBAM without periplasmic density). For PgBAM with periplasmic density, after the 2 initial rounds of 2D classification following template picking, the particles were subjected to further 5 rounds of 2D classification (Supplementary Fig. 3). The resulting 13,929 particles were subjected to heterogeneous refinement with one protein and 2 decoy volumes. Protein class was used in non-uniform refinement, yielding a final reconstruction at 4.26 Å (PgBAM with periplasmic density). Global resolution was estimated using gold-standard Fourier shell correlation curves and the 0.143 criterion (Supplementary Fig. 3c). Particle viewing direction distribution plots and local resolution estimates are displayed in Supplementary Fig. 3d-f. AF2 models of all subunits were docked into the final volume of PgBAM without periplasmic density and subjected to cycles of manual adjustment in Coot and real space refinement in Phenix. Mannose monosaccharides were placed in O-glycan densities to prevent the protein model from moving into the glycan density. Model refinement statistics are shown in Supplementary Table 1.

### Growth curves

Overnight EBHI cultures of *B. theta* were used to inoculate 5 ml of fresh EBHI (1:25), followed by incubation at 37°C for 3-4 h in an anaerobic cabinet. The cells were collected by centrifugation for 3 min at 2,900*g*, 20°C, and resuspended in 1 ml pre-warmed MM. Glass tubes with EBHI or MM (including carbon source) and supplements indicated in the main text were inoculated with the resuspended cells at OD_600_=0.04, and the cultures were dispensed in 0.2 ml aliquots in sterile 96-well clear polystyrene cell culture plates (Corning Costar). Medium without any added cells was included as blank. Plates were incubated anaerobically at 37°C for 30 min and sealed with a Breathe-Easy gas-permeable seal (Diversified Biotech), followed by growth monitoring for 48 h in a Biotek Epoch microplate reader housed inside an anaerobic cabinet. Blank well readings were subtracted from wells with added cells, and triplicate readings were averaged. The experiments were repeated at least two times on different days to ensure consistency.

*P. gingivalis ΔbamG, ΔbamH and ΔbamJ* mutants were first grown in eTSB overnight. The cultures were resuspended in fresh eTSB or washed and resuspended in DMEM supplemented with 1% bovine serum albumin, 0.25 g/l L-cysteine and either regular or decreased amounts of vitamin K (0.5 mg/l and 0.05 mg/ml, respectively) and hemin (5 mg/ml and 0.5 mg/ml, respectively). In each case the OD_600_ was adjusted to 0.2 and the growth was monitored for 48 h through OD_600_ measurements at predetermined time intervals, since *P. gingivalis* does not grow in multi-well plates.

### PgBAM expression and purification

The PgBAM complex was isolated from His-tagged BamG-expressing *P. gingivalis* according to a previously described protocol^42^. Briefly, cells from 5 L were harvested by centrifugation at 6,500 g for 40 min and resuspended in TBS with 1 mM *N*_α_-tosyl-L-lysine chloromethyl ketone (TLCK), 1× cOmplete EDTA-free Protease Inhibitor Cocktail and 20 μg/ml DNase I, followed by lysis with Constant Systems Cell Disruptor at 23 kpsi. The total membrane fraction was separated from the soluble fraction through ultracentrifugation at 200,000 g for 1 h and then homogenized and solubilized in TBS with 1% n-dodecyl β-D-maltoside (DDM) at 4°C overnight. The extract was clarified by ultracentrifugation at 200,000 g for 50 min and loaded on 1.5 ml pre-equilibrated nickel-affinity resin (Chelating Sepharose). The column was washed with 15 CV of TBS containing 0.2% DDM and 30 mM imidazole, and the bound protein was eluted with 2 CV of TBS containing 0.2% DDM and 250 mM imidazole. The complex was further purified by size exclusion chromatography in 10 mM HEPES pH 8.0, 100 mM NaCl, 0.03% DDM using a Superdex 200 Increase 10/300 GL column. In pulldown experiments shown in Fig. 5g,h, RagB-His from wild-type and *ΔbamG* cells was purified in the same manner, except lauryldimethylamine-*N*-oxide (LDAO) was used instead of DDM.

### Pulldown proteomics sample preparation, mass spectrometry and analysis

Protein solutions were diluted 1:1 with aqueous 10% (v/v) sodium dodecyl sulfate, 100 mM triethylammonium bicarbonate (TEAB). Protein was reduced with 5.7 mM tris(2-carboxyethyl)phosphine and heating to 55°C for 15 mins before alkylation with 22.7 mM methyl methanethiosulfonate at room temperature for 10 mins. Protein was acidified with 6.5 μL of aqueous 27.5 % (v/v) phosphoric acid then precipitated with dilution seven-fold into 100 mM TEAB 90% (v/v) methanol. Precipitated protein was captured on S-trap (Profiti – C0-micro) and washed five times with 165 µL 100 mM TEAB 90% (v/v) methanol before digesting with the addition of 20 µL 0.1 µg/µL Promega Trypsin/Lys-C mix (V5071) in aqueous 50 mM TEAB and incubation at 47 °C on hot plate for 2h. Peptides were recovered from S-trap by spinning at 4000 g for 60 s. S-traps were washed with 40 µL aqueous 0.2% (v/v) formic acid and 40 µL 50% (v/v) acetonitrile:water and washes combined with the first peptide elution. Peptide solutions were dried in a vacuum concentrator then resuspended in 20 μL aqueous 0.1% (v/v) formic acid. Peptides were loaded onto EvoTip Pure tips for desalting and as a disposable trap column for nanoUPLC using an EvoSep One system. A pre-set EvoSep 100 SPD gradient (from Evosep One HyStar Driver 2.3.57.0) was used with an 8 cm EvoSep C_18_ Performance column (8 cm x 150 μm x 1.5 μm). The nanoUPLC system was interfaced to a timsTOF HT mass spectrometer (Bruker) with a CaptiveSpray ionisation source (Source). Positive PASEF-DDA, ESI-MS and MS^2^ spectra were acquired using Compass HyStar software (version 6.2, Bruker). Instrument source settings were: capillary voltage, 1,500 V; dry gas, 3 l/min; dry temperature; 180°C. Spectra were acquired between *m/z* 100-1,700. TIMS settings were: 1/K0 0.6-1.60 V.s/cm^2^; Ramp time, 100 ms; Ramp rate 9.42 Hz. Data dependant acquisition was performed with 10 PASEF ramps and a total cycle time of 1.17 s. An intensity threshold of 2,500 and a target intensity of 20,000 were set with active exclusion applied for 0.4 min post precursor selection. Collision energy was interpolated between 20 eV at 0.6 V.s/cm^2^ to 59 eV at 1.6 V.s/cm^2^.

LC-MS data, in Bruker .d format, was processed using DIA-NN (1.8.2.27) software and searched against an in-silico predicted spectral library, derived from the *Bacteroides thetaiotaomicron* subset of UniProt (UP000001414, 2024/06/10) appended with common proteomic contaminants. Search criteria were set to maintain a false discovery rate (FDR) of 1%. High-precision quant-UMS^77^ was used for extraction of quantitative values within DIA-NN. Peptide-centric output in .tsv format, was pivoted to protein-centric summaries using KNIME 5.1.2 and data filtered to require protein q-values < 0.01 and a minimum of two peptides per accepted protein. Calculation of log2 fold difference and differential abundance testing was performed using limma via FragPipe-Analyst^78^. Sample minimum imputation was applied and the Hochberg and Benjamini approach was used for multiple test correction. Volcano plots were made using the EnhancedVolcano^79^ package in R.

### Quantitative proteomics sample preparation, mass spectrometry, and data analysis

For *B. theta* membrane proteomics, three independent 0.5 l cultures wild type and *ΔbamG* strains were grown and processed up to the membrane isolation step as described above for BtBAM complex purification. For *P. gingivalis* whole cell proteomics, three independent wild type and *ΔbamG P. gingivalis* cultures were grown overnight in eTSB. Cells from 50 ml of culture were washed three times with ice-cold PBS. Aliquots corresponding to 50 ml culture of OD_600_ = 0.6 were centrifuged and the pellets were stored at −80°C until needed.

*P. gingivalis* and *B. theta* cell and membrane pellets were lysed by addition of 50 mM triethyl ammonium bicarbonate (TEAB) pH 8.5 with 5 % (w/v) SDS and sonication using a UP200St ultrasonic processor (Hielscher) at 90 W, 45 s pulse, 15 s rest, three times. *B. theta* lysis buffer included 1x cOmplete protease inhibitor cocktail (Roche), for *P. gingivalis*, 2x protease inhibitor cocktail and 100 mM N-ethylmaleimide (NEM). Samples were then denatured with 5 mM tris(2-carboxyethyl)phosphine (TCEP) at 60 °C for 15 minutes, alkylated with 30 mM iodoacetamide (*B. theta*) or 10 mM NEM (*P. gingivalis*) at room temperature for 30 minutes in the dark, and acidified to a final concentration of 2.7 % phosphoric acid. Samples were then diluted eightfold with 90 % MeOH 10 % TEAB (pH 7.2) and added to the S-trap (Protifi) micro columns. The manufacturer-provided protocol was then followed, with a total of five washes in 90 % MeOH 10 % TEAB (pH 7.2), and trypsin added at a ratio of 1:10 enzyme:protein and digestion performed for 18 h at 37 °C. Peptides were dried using a vacuum concentrator and stored at −80 °C, and immediately before mass spectrometry were resuspended in 0.1 % formic acid.

Liquid chromatography (LC) was performed using an Evosep One system with a 15 cm Aurora Elite C18 column with integrated captivespray emitter (IonOpticks), at 50 °C. Buffer A was 0.1 % formic acid in HPLC water, buffer B was 0.1 % formic acid in acetonitrile. Immediately prior to LC-MS, peptides were resuspended in buffer A and a volume of peptides equivalent to 500 ng was loaded onto the LC system-specific C18 EvoTips, according to manufacturer instructions, and subjected to the predefined Whisper100 20 SPD protocol (where the gradient is 0-35 % buffer B, 100 nl/min, for 58 minutes; 20 samples per day are permitted using this method). The Evosep One was used in line with a timsToF-HT mass spectrometer (Bruker), operated in diaPASEF mode. Mass and IM ranges were 300-1200 *m/z* and 0.6-1.4 1/*K_0_*, diaPASEF was performed using variable width IM-*m/z* windows, as described previously^80^. TIMS ramp and accumulation times were 100 ms, total cycle time was ∼1.8 seconds. Collision energy was applied in a linear fashion, where ion mobility = 0.6-1.6 1/K_0_, and collision energy = 20 - 59 eV.

Raw diaPASEF data files were searched using DIA-NN V 1.8.2 beta 27^81^, using its *in silico* generated spectral library function, based on reference proteome FASTA files for *B. thetaiotaomicron* (UP000001414, downloaded from UniProt on 01/12/2021) or *P. gingivalis* (UP000008842, downloaded from UniProt on 13/02/2024), and a common contaminants list^82^. Trypsin specificity with a maximum of 2 variable modifications and 1 missed cleavage were permitted per peptide, cysteine carbamidomethylation (*B. theta*) or NEM (UniMod:108, *P. gingivalis*) were set as fixed, oxidation of methionine was set as a variable modification. Peptide length and m/z was 7-30 and 300-1200, charge states 2-4 were included. Mass accuracy was fixed to 15 ppm for MS1 and MS2. Protein and peptide FDR were both set to 1 %. Match between runs and RT normalisation were used for *B. theta* but not *P. gingivalis*. All other settings were left as defaults. The protein group matrix outputs were processed (separately for each species) in R using Rstudio, with filtering to exclude contaminants, and include a minimum of 2 peptides per protein group, and proteins present in a minimum of 2/3 of replicates in any one condition. Statistical analysis was performed using Limma^83^, with statistical significance inferred by T-test with Benjamini Hochberg correction for multiple comparisons. Volcano plots were made using the EnhancedVolcano^79^ package in R.

### *B. theta* proteinase K shaving assay

Overnight EBHI cultures were used to inoculate fresh minimal medium supplemented with 0.5% fructose, or 0.5% maltose for the control *susA_his_* strain, at OD_600_∼0.1. After a 4-hour incubation at 37°C in an anaerobic chamber, 33 OD units of each strain were harvested by centrifugation for 5 min at 4,000*g*. Cell pellets were resuspended in 1 ml PK buffer (50 mM Tris-HCl pH 8.0, 50 mM NaCl, 1 mM MgCl_2_), split into four 250 μl aliquots, and pelleted again. Two aliquots were resuspended in 180 μl PK buffer and supplemented with 20 μl PK buffer or 20 mg/ml fresh proteinase K solution. The other two aliquots were resuspended in 180 μl PK buffer containing 1% Triton X-100 (v/v) and supplemented with 20 μl PK buffer or 20 mg/ml fresh proteinase K solution. Proteinase K digestion was carried out at 37°C for 2 h under aerobic conditions. Samples containing Triton X-100 were supplemented with 5 mM phenylmethylsulfonyl fluoride (PMSF) and incubated at 20°C for 15 min. Samples without detergent were pelleted, washed twice with PK buffer containing 2 mM PMSF, and lysed in 200 μl PK buffer with 1% Triton X-100 and 2 mM PMSF for 10 min at 20°C. Samples were mixed with SDS loading buffer, boiled immediately for 10 min, and separated on a 12% FastCast gel (Bio-Rad). Proteins were transferred from the gel to a 0.2 μm polyvinylidene fluoride membrane using a Bio-Rad Trans-Blot Turbo system. The membrane was stained with Ponceau S to confirm successful transfer, blocked for 1 hour with 3% BSA solution in PBST (phosphate buffered saline with 0.1% Tween-20), and incubated for 2 h with anti-His-HRP conjugate monoclonal antibody (Roche) diluted 1:500 in blocking solution. The membrane was washed three times with PBST and HRP was detected using chemiluminescence. The membrane was then re-probed with StrepTactin-HRP (Bio-Rad) in blocking solution (1:10,000) for 2 h, and washed and developed as above.

### *P. gingivalis* cell fractionation

*P. gingivalis* strains were grown overnight in eTSB. All cultures were adjusted to the same OD_600_ and centrifuged at 7,500 g for 40 min. The supernatant was further ultracentrifuged at 200,000 g for 1 h to separate outer membrane vesicles (OMVs) from media. OMV-free media were concentrated 20×. OMVs were resuspended in PBS with 1 mM TLCK and sonicated. Cells were washed and resuspended in PBS with 1 mM TLCK, followed by sonication and centrifugation at 7,500 g for 15 min. Supernatant samples were collected and processed as for the soluble fraction. Pellets, corresponding to the insoluble fraction, were washed and resuspended in PBS with 1 mM TLCK.

### SDS-PAGE sample preparation and western blotting analysis

Samples of *P. gingivalis* OMVs, OMV-free media, soluble and insoluble fractions were mixed with NuPAGE LDS Sample Buffer and boiled at 95°C for 5 min. Dithiothreitol was added to 50 mM and the samples were boiled again at 95°C for 5 min. Samples were separated in NuPAGE Bis-Tris Mini Protein Gels, 4–12% and either stained with Coomassie Brilliant Blue G250 or electrotransferred onto nitrocellulose membranes in 25 mM Tris, 192 mM glycine and 20% methanol at 100 V for 60 min. To visualize total protein, membranes were stained with 0.1% PonceauS in 1% acetic acid and rinsed with distilled water. Membranes were blocked with 5% skim milk (SM) in TBS with 0.1% Tween-20 (TTBS) at 4°C overnight followed by 1 h incubation with anti-HmuY antibodies and 50 min incubation with secondary HRP-conjugated anti-rabbit antibodies at room temperature. Signal detection was carried out with Pierce ECL Western blotting substrate.

### Transmission electron microscopy with sectioned *B. theta* cells

Cells were cultured to stationary phase (OD_600_ = 1.5 – 1.8), harvested by centrifugation (5,000 × g, 5 min) and washed twice with 1 mM HEPES pH 7.0, 55 mM sucrose. Samples were fixed in 1.5% (v/v) glutaraldehyde, 0.5% OsO_4_, 0.15% ruthenium red, 55 mM sucrose in 1 mM HEPES pH 7.0. Next, samples were dehydrated in an ethanol series (50%, 70%, 90%, 96%, 100%, 2 × 15 min each) and in propylene oxide (2 × 10 min) before embedding in a mixture of epoxy resins (Poly/bed 812, MNA, DDSA) (Polysciences). The following day the preparations were transferred to DMP-30 resin and polymerized in embedding forms at 60°C. Embedded samples were sectioned with an EM UC7 microtome (Leica). Sections were contrasted on the grids using uranyl acetate and lead citrate. A Tecnai Osiris electron microscope (FEI) operating at an accelerating voltage of 200 kV was used to image the stained sections. Gatan Microscopy Suite (GMS) software v3.4.3 was used to take images on a Rio16 camera (Gatan).

### Bioinformatics

To determine the taxonomic distribution of newly discovered BAM components, protein sequences were first subjected to SSDB Motif Search in KEGG database. Then, the conserved motif was used as a query in the Enzyme Function Initiative - Enzyme Similarity Tool^84^. Default parameters and the UniProt database were used for searches.

Homologues to BamF (Bt4367) were identified through complementary BlastP searches performed at the NCBI Bast server against the non-redundant protein sequences (nr)^85^. Bt4367 and selected BamF homologues were used as queries in BlastP unrestricted and taxonomically restricted searches to identify a representative set of BamF homologues across sequence and taxonomic diversity. A selection of BamF homologues were combined to generate protein alignments, optimising taxonomic and sequence diversity representation for phylogenetic inferences. SeaView v5.0.5^86^ was used to infer, manipulate, and visualise the alignments to generate figures and initial phylogenies. Clustal Omega was used to generate the protein alignments^87^ and Gblocks^88^ was used to trim the alignments to select the sites with the strongest hypothesis of homology and to remove excessively divergent regions prior to phylogenetic inference. The maximum likelihood framework for protein phylogenetics implemented in IQ-TREE was used for tree inferences^89,90^. Support values for branches were estimated using ultrafast bootstrapping^91^. The best fitting single-matrix homogenous substitution model for the alignment was inferred with the automatic model selection (function “Auto”), which was LG+G4+I+F (Optimal pinv=0.045, alpha=1.879) for the Akaike information (AIC) and corrected AIC criterion and the Bayesian information criterion (BIC).

To investigate the distribution of BamF homologues across Bacteroidetes genomes, the annotated proteins from complete genomes from the RefSeq genome database at the NCBI^92^ were downloaded (1448 genomes on 17 September 2024). A local BLASTP 2.12.0+ search^93,94^ was performed using Bt4367/BamF as query (*Bacteroides thetaiotaomicron* VPI-5482, accession AAO79472.1 putative outer membrane protein, Length=431 residues). Blast hits with E-value ≤ 0.05 were recorded and are listed in Supplementary Table 2. tBlastn searches using the NCBI Blast portal were performed against individual genomes without strong BlastP hits (Supplementary Discussion).

## Author contributions

AS determined cryo-EM structures, performed experiments in *B. theta*, generated figures and wrote the paper. MM determined cryo-EM structures, generated figures and wrote the paper and performed experiments in *P.gingivalis* together with KM. AMF carried out proteomics, supervised by MT. RPH and AJH carried out phylogenetic analyses, with AJH supervised by RPH. AB managed the Newcastle University Structural Biology Facility. CS and JJE carried out proteomics. BvdB conceived the project, performed experiments in *B. theta* and wrote the paper.

## Data availability

Pulldown proteomic mass spectrometry data sets and results files are referenced in ProteomeXchange (PXD058163) and available to download from MassIVE (MSV000096498) [doi:10.25345/C57W67J00]. Pre-publication access can be obtained with the following link ftp://MSV000096498@massive.ucsd.edu (username MSV000096498, password S73xCTORvm9SM2m7). *B. theta* membrane and *P. gingivalis* whole cell mass spectrometry proteomics data have been deposited to the ProteomeXchange Consortium via the PRIDE^95^ partner repository with the dataset identifiers PXD058903 for *B. theta* and PXD058905 for *P. ginigivalis.* Reviewers can access the datasets by logging in to the PRIDE website using the following account details for *B. theta*: username, password afLrMXZevbBq; or *P. gingivalis*: project accession PXD058905, token: eAL4YgXN00lJ. Cryo-EM reconstructions and atomic coordinate files with the indicated accession numbers have been uploaded to the Electron Microscopy Data Bank (EMDB) and the Protein Data Bank (PDB): BtBAM class 1 (EMD-52200 and 9HIS), BtBAM class 2 (EMD-52202 and 9HIV), BtBAM composite map and model (EMD-52209 and 9HJ3), PgBAM class 1 (EMD-52218 and 9HJM), and PgBAM class 2 (EMD-52219).

## Supporting information

Supplementary Discussion

## Acknowledgements

We thank Michael Fischbach (Stanford University) for providing the Bt1927-ON strain and Javier Abellon-Ruiz (Newcastle University) for providing the *B. theta* SusA_his_ strain. BvdB was financially supported by a Wellcome Trust Investigator award (214222/Z/18/Z), providing salary support to AS. The York Centre of Excellence in Mass Spectrometry was created thanks to a major capital investment through Science City York, supported by Yorkshire Forward with funds from the Northern Way Initiative, and subsequent support from EPSRC (EP/K039660/1; EP/M028127/1). We acknowledge the use of cryo-EM facilities at the Astbury Biostructure Laboratory, which was financially supported by the University of Leeds and the Wellcome Trust (108466/Z/15/Z and 221524/Z/20/Z), and thank Yehuda Halfon for support. This study was supported by a Polish National Science Centre grant to MM (UMO-2023/51/D/NZ1/02675). MM was also supported by the Bekker programme of the Polish National Agency for Academic Exchange (PPN/BEK/2020/1/00103/U/00001). We acknowledge the Polish high-performance computing infrastructure PLGrid (HPC Center: ACK Cyfronet AGH) for access to computational facilities and support (PLG/2023/016774). This publication was partially developed under the provision of the Polish Ministry and Higher Education project “Support for research and development with the use of research infrastructure of the National Synchrotron Radiation Centre SOLARIS” under contract nr 1/SOL/2021/2 (project ID 233012, cryo-electron microscope). We thank Michał Rawski, Grzegorz Ważny, Marcin Jaciuk and Paulina Indyka (all from SOLARIS Centre) for assistance in preparation of grids and imaging. We thank Jan Potempa (Jagiellonian University) for valuable discussions during the project and for financial support providing salary to MM and covering costs of materials and services from his financial resources (NCN grant UMO-2018/30/A/NZ5/00650 and others at the Jagiellonian University. Barbara Potempa (University of Louisville) is acknowledged for technical support. We thank Teresa Olczak (University of Wroclaw) for providing anti-HmuY antibodies. We thank Olga Barczyk-Woznicka (Jagiellonian University) and Michal Pacia (Jagiellonian University) for assistance with TEM experiments.

## Extended Data Figures

**Extended Data Figure 1.**
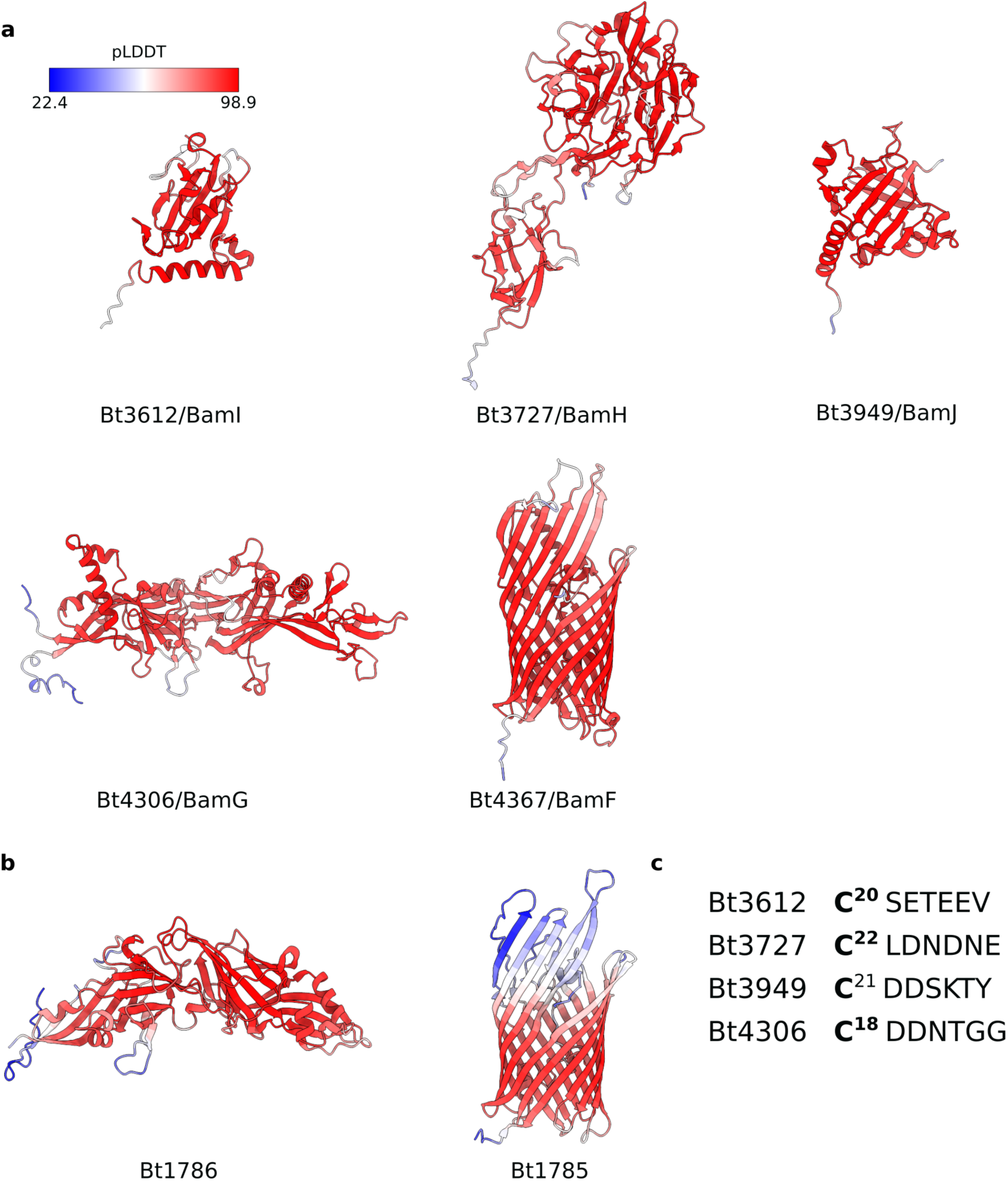
Structure predictions of proteins of interest. **a**, AF2^28^ predictions of abundant proteins identified in BtBamA_his_ pulldowns. Protein models are coloured by pLDDT values (colour key). **b**, AF2 predictions of Bt1786 and Bt1785, which are homologues of Bt4306/BamG and Bt4367/BamF, respectively. Only the mature part of each protein is shown. **c**, The lipid anchor cysteine and six subsequent amino acid residues of each lipoprotein detected in BtBamA_his_ pulldowns. The presence of two or more acidic residues indicates that these lipoproteins have a lipoprotein export signal and are likely surface exposed^24,25^.

**Extended Data Figure 2.**
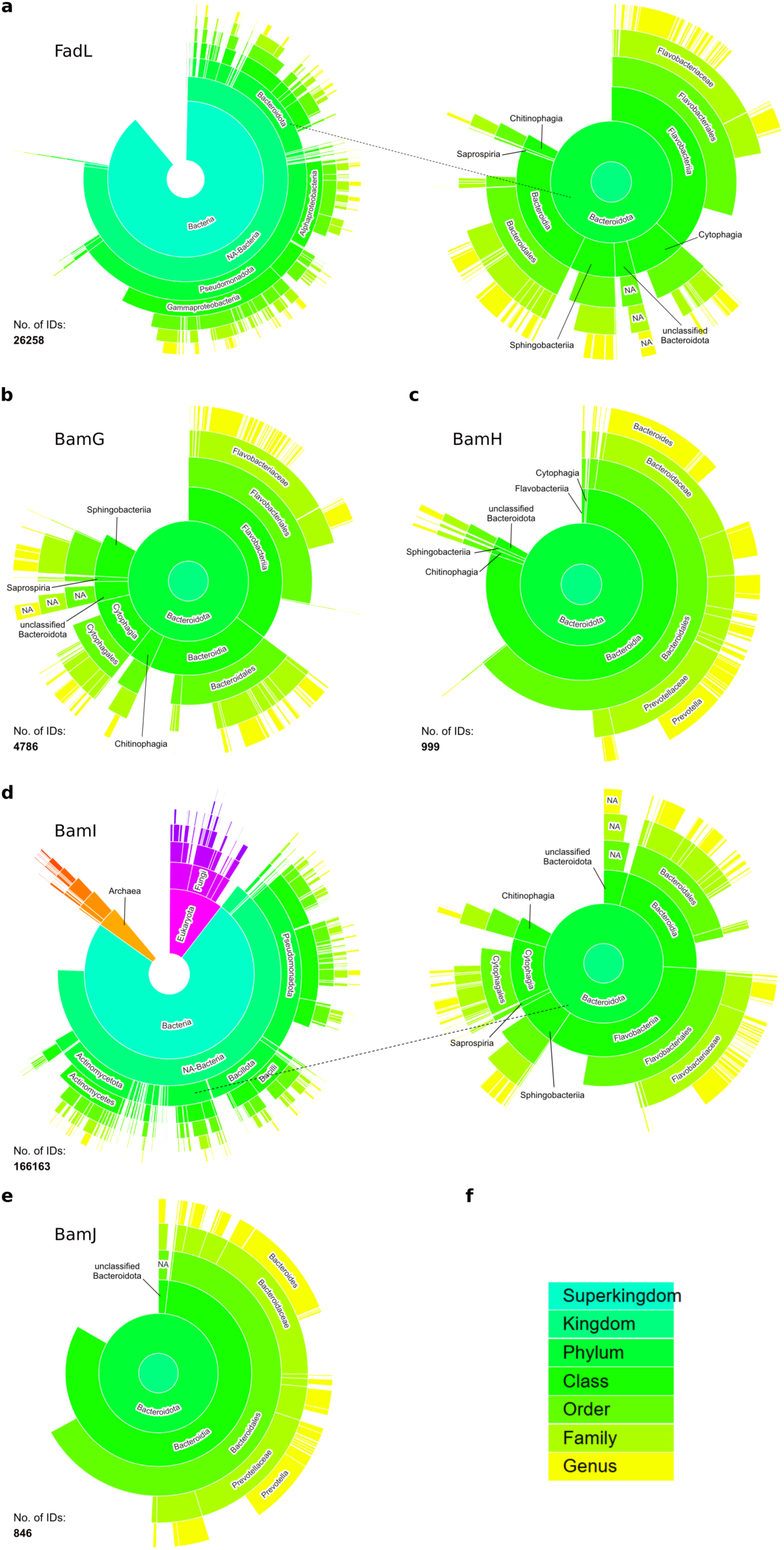
Distribution of BAM-associated surface-exposed lipoproteins in Bacteria. Organisms containing homologues of Bt4367/FadL/ (**a**), Bt4306/BamG (**b**), Bt3727/BamH (**c**), BamI/Bt3612 (**d**) and BamJ/Bt3949 (**e**). **f**, Taxonomic level colour key. Displayed number of IDs corresponds to the number of pBLAST hits. The taxonomy sunburst plots were generated using the Enzyme Function Initiative - Enzyme Similarity Tool^84^. NA, unclassified.

**Extended Data Figure 3.**
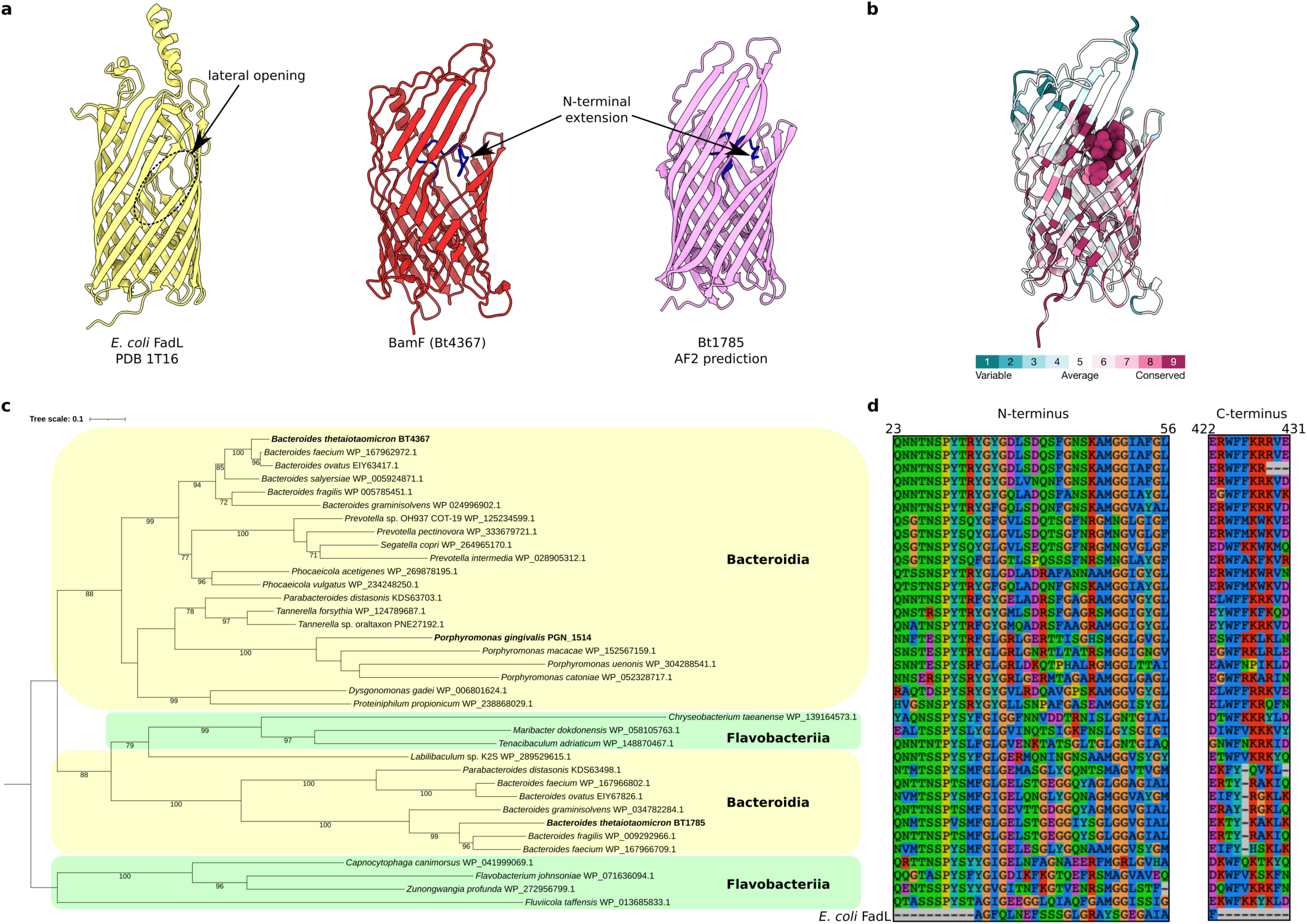
Properties and phylogenetics analysis of BamF. **a**, Comparison of the *E. coli* FadL crystal structure (PDB 1T16^30^), the BtBamF cryo-EM structure and the AF2 prediction of Bt1785 (signal sequence not shown). Views generated from a superposition. The 12-amino acid N-terminal extension of BtBamF and Bt1785 is shown in blue. **b**, ConSurf analysis of BtBamF. The conserved N-terminal extension (residues 23-37) is shown in sphere representation. **c**, BamF protein phylogeny focusing on homologues encoded by selected members of the Bacteroidia and Flavobacteriia classes. The phylogeny was inferred from an alignment of 240 residues, and it is routed with a selection of members of the class Flavobacteriia, the closest outgroup to the Bacteroidia identified by phylogenomics^96^. The phylogeny was rooted with the lineage that includes the most basal branch among the Flavobacteriia, *Fluviicola taffensis*^96^. However, the phylogeny inferred from a broader taxa sampling does not resolve the position of the two distinct Flavobacteriia lineages (highlighted in green). The non-monophyly of the genus *Bacteroides* among the Bacteroidia (highlighted in yellow) and Flavobacteriia could be due to a phylogenetic inference artefact or reflect a complex evolution of the BamF genes, including potential lateral gene transfers and/or gene duplications. Ultrafast bootstrap values (1000 replicates) below 95% indicate a lack of resolution for some of the branches to support either possibility. The strongly supported distinct positions for the two BamF homologues encoded by the species *Parabacteroides distasonis* (Family Tannerellaceae) suggest a lateral gene transfer from a *Bacteroides* species into *P. distasonis* (accession KDS63498.1). The scale bar indicates the inferred mutation per aligned residues for the automatically selected model (LG+G4+I+F). The accession number for each protein in the alignment is indicated. **d**, Alignments of the N- and C-terminal regions of Bacteroidetes BamF homologues showing conserved terminal extensions compared to *E. coli* FadL. Rows correspond to the sequences shown in the phylogeny in **c**; the bottom row shows the *E. coli* FadL sequence. Residue numbers at the top correspond to the Bt4367/BamF sequence.

**Extended Data Figure 4.**
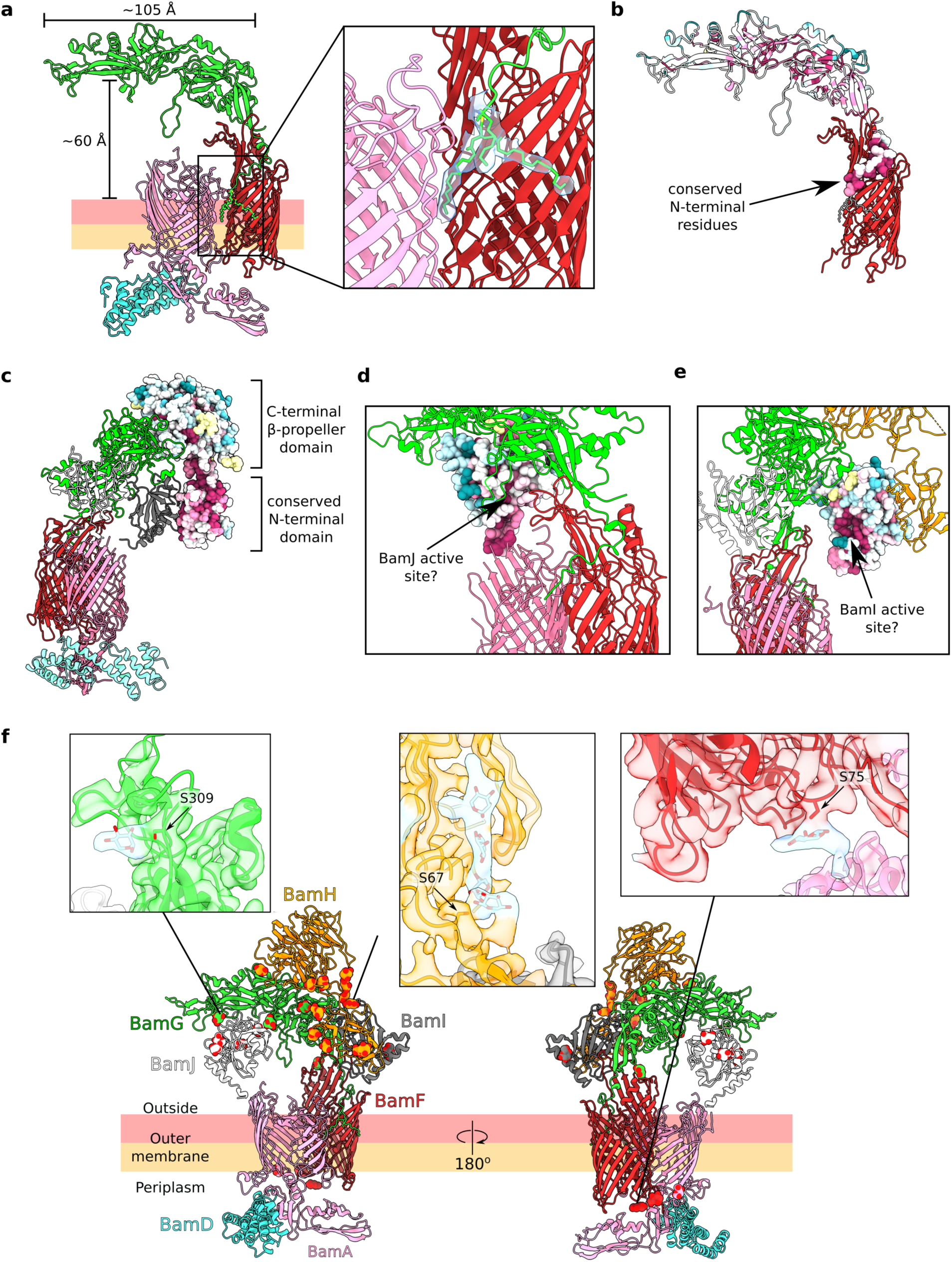
Properties and conservation analysis of novel BtBAM components. **a**, BtBamG extends in an arch above the outer membrane, with its N-terminal cysteine lipid anchor wedged between BtBamA and BtBamF β-barrels. The inset shows the map-model fit of the lipid anchor. **b-e**, ConSurf^35^ analyses of BtBamG (**b**), BtBamH (**c**), BtBamI (**d**) and BtBamJ (**e**), using default program parameters (minimum and maximum sequence identity set to 35% and 95%, respectively). Conserved BtBamG N-terminal residues 18-34 are shown in sphere representation in **b**. Putative BamI and BamJ PPI active sites face the interior of the extracellular dome in **d** and **e**. **f**, O-glycosylation sites displayed as mannose monosaccharides in space filling representation. The mannose residues were placed into the glycan densities to stabilise protein model refinement. The exact chemical nature of the *B. theta* O-glycan is unknown. Insets show three representative glycans in blue extending from BamG Ser309, BamH Ser67, and BamF Ser75 sidechains.

**Extended Data Figure 5.**
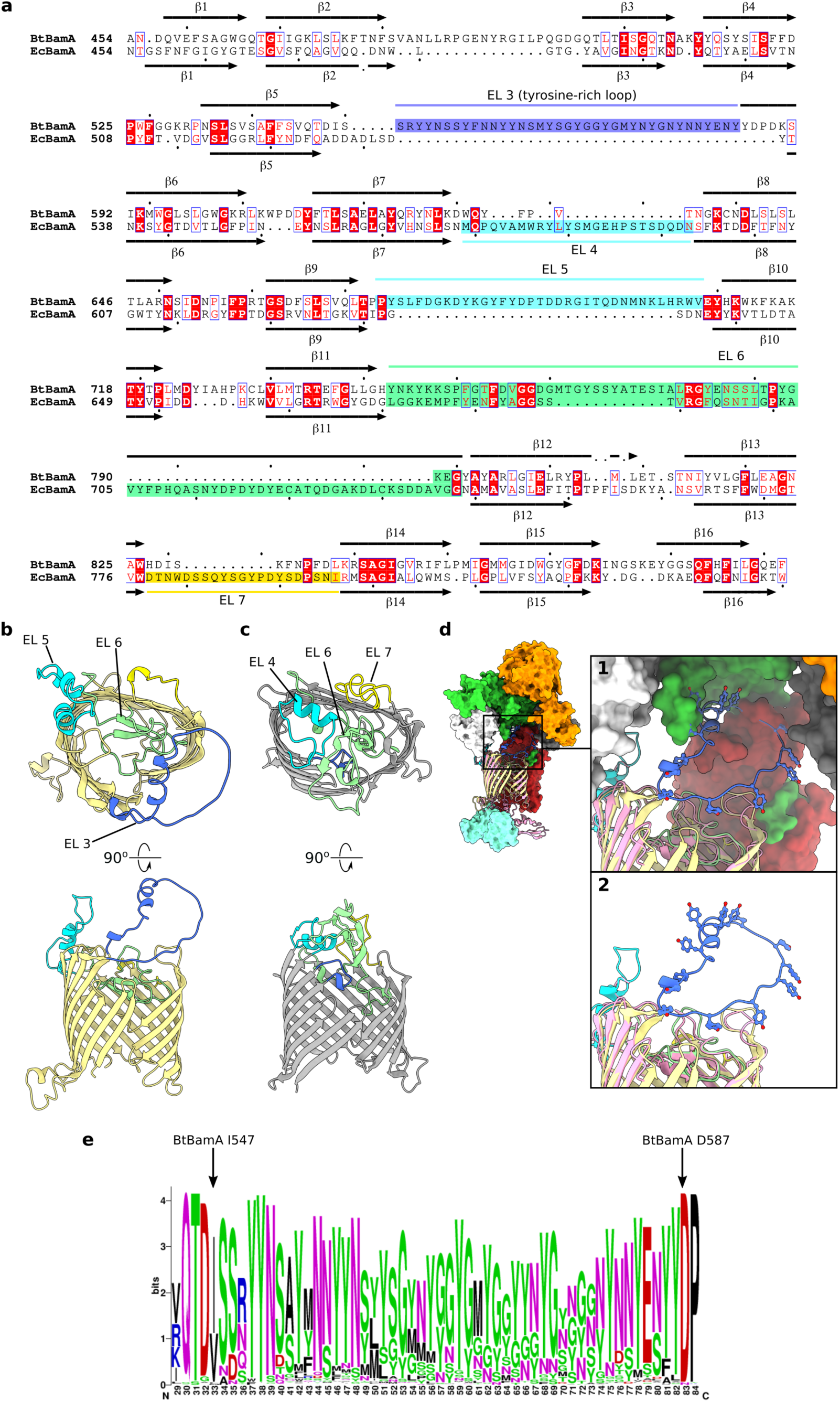
Comparison of *B. theta* and *E. coli* BamA β-barrels. **a**, Structure-based sequence alignment of BtBamA and *E. coli* BamA (EcBamA). Sequence identity – 22%, TM-align score – 0.64. The β-barrel strands of BtBamA and EcBamA are annotated, respectively, above and below the sequence alignment. Extracellular loops (EL) of interest are annotated. **b**, **c**, AF2 models of BtBamA (**b**) and EcBamA (**c**) β-barrels with extracellular loops coloured as in (**a)**. **d**, The position of the tyrosine-rich loop (EL 3) in the context of the whole BtBAM complex. The AF2 model of BtBamA, coloured as in **b**, is superposed on the experimental BtBamA structure. No cryo-EM density for the tyrosine-rich loop was observed. Inset 1 shows a close-up view of EL3 inside the dome. Inset 2 shows the same view of BtBamA as inset 1 with the other BAM components omitted for clarity. **e**, WebLogo^97^ representation of the Bacteroidetes BamA EL3 region generated from 861 aligned BlastP hits shows enrichment of tyrosine, glycine and asparagine residues.

**Extended Data Figure 6.**
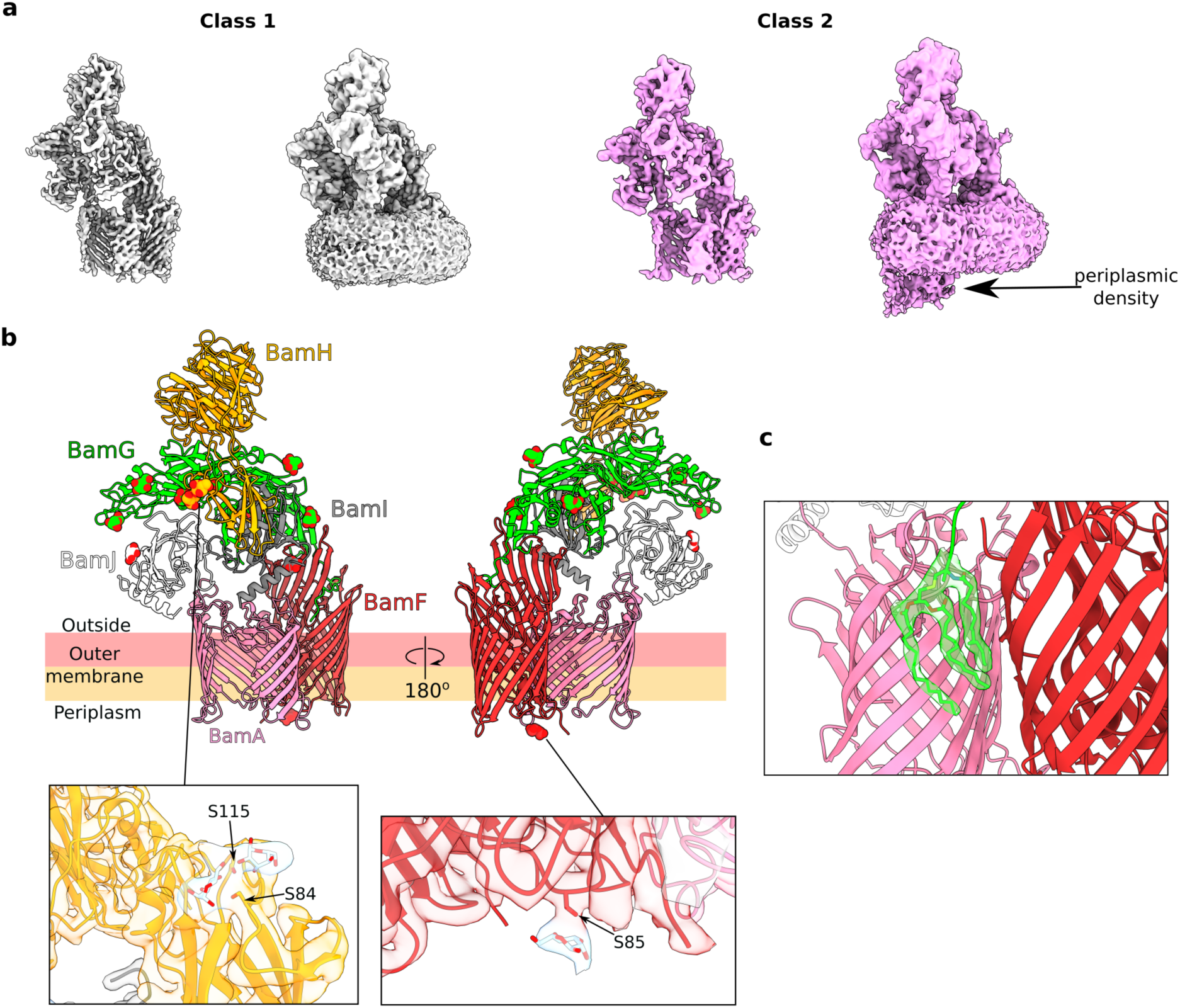
PgBAM structural features. **a**, Two cryo-EM classes of PgBAM shown at high (left) and low (right) contour. The periplasmic density observed in class 2 is assumed to correspond to PgBamD and/or PgBamA POTRA domain 5. **b**, Glycosylation sites observed in the PgBAM structure. Mannose units (space filling representation) were built into the glycan density to stabilise protein model refinement. Insets show close-ups of representative model density at glycosylation sites on BamH (Ser84, Ser115) and BamF (Ser85). **c**, Model-density fit of the PgBamG lipid anchor at the PgBamAF barrel interface.

**Extended Data Figure 7.**
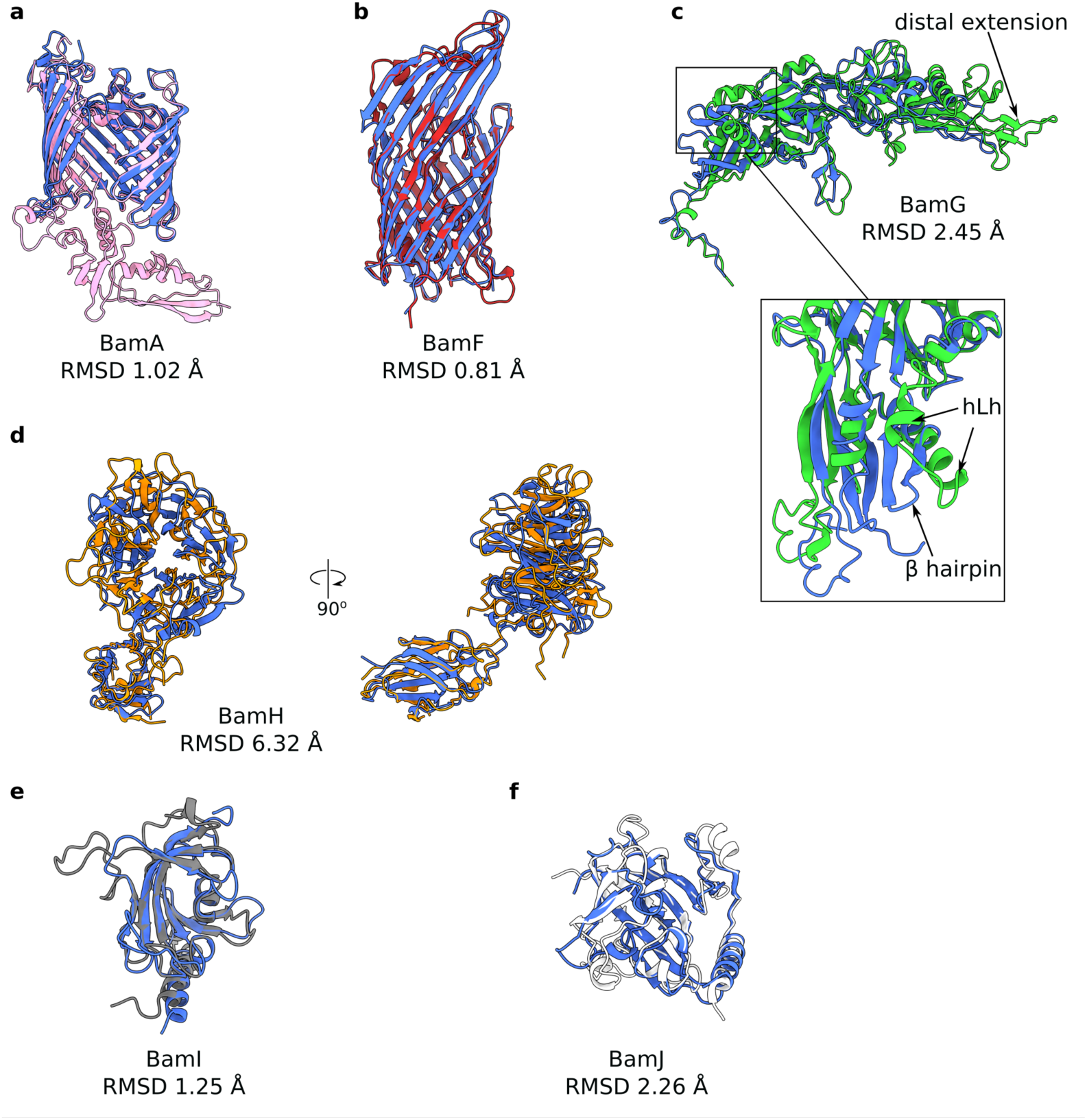
Structural alignment of individual BtBAM and PgBAM components. Models were superimposed using Matchmaker in ChimeraX^98^, and the Cα-Cα RMSD between pruned atom pairs is reported. BtBAM components are coloured using the same colour scheme as in Figure 2 of the main text; PgBAM components are shown in blue. hLh, helix-loop-helix.

**Extended Data Figure 8.**
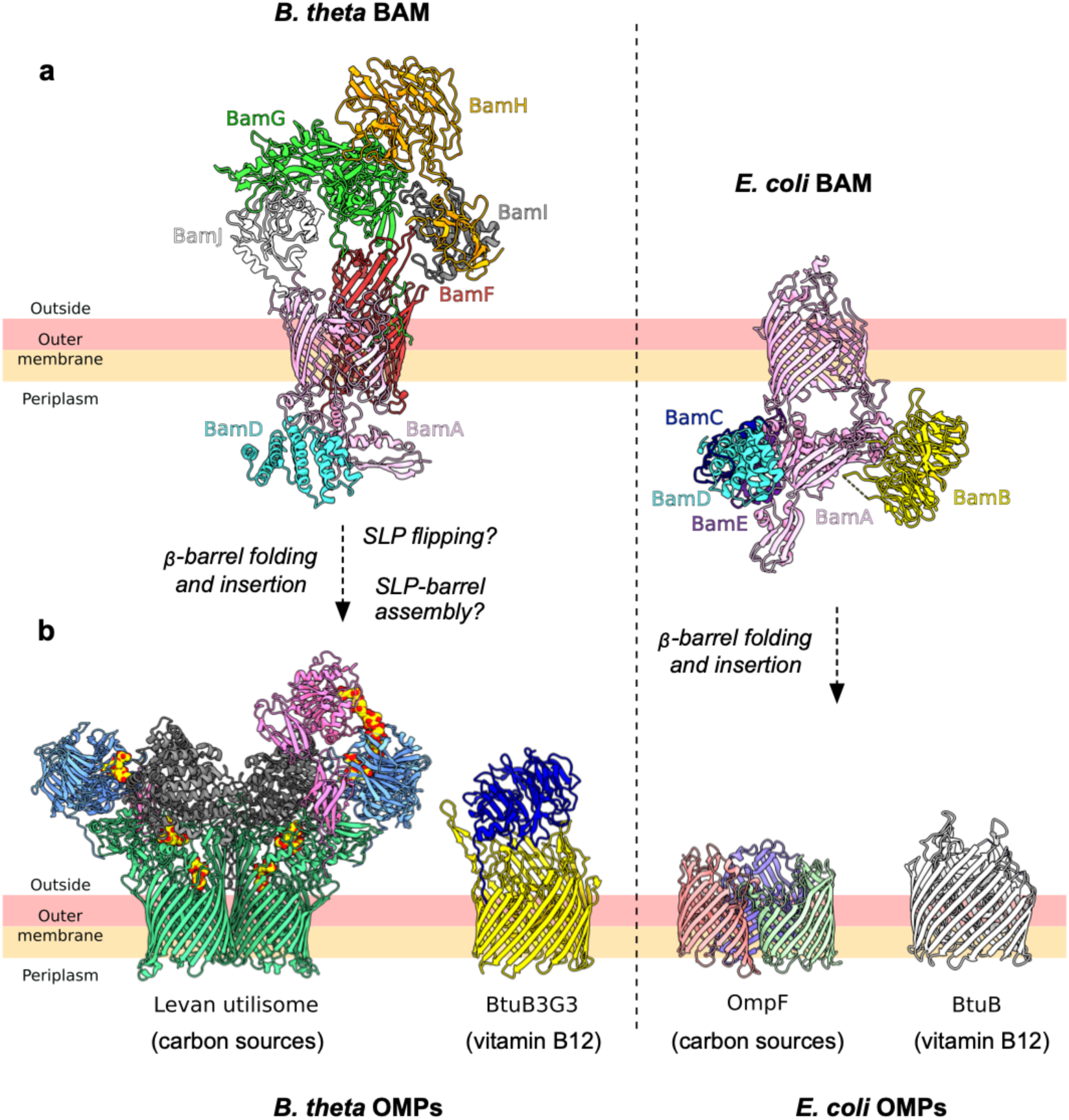
Comparison of *B. theta* and *E. coli* BAM complexes and their typical substrates. **a**, Comparison of BtBAM (PDB 9HJ3) and EcBAM (PDB 5D0O^3^) structures. The views were generated from a superposition of the BamA chains. **b**, Examples of *B. theta* and *E. coli* BAM substrates: levan utilisome (Bt1760-3), PDB 8AA2^46^; vitamin B12 transporter BtuB3G3 (Bt2094-5), PDB 8P98^99^; vitamin B12 transporter BtuB, PDB 1NQE^100^; general porin OmpF, PDB 1OPF^101^. As is clear from the comparison of OM transporters for carbon sources and vitamin B12, the Bacteroidetes BAM complex likely requires expanded functionality compared to its Proteobacterial counterpart.

**Extended Data Figure 9.**
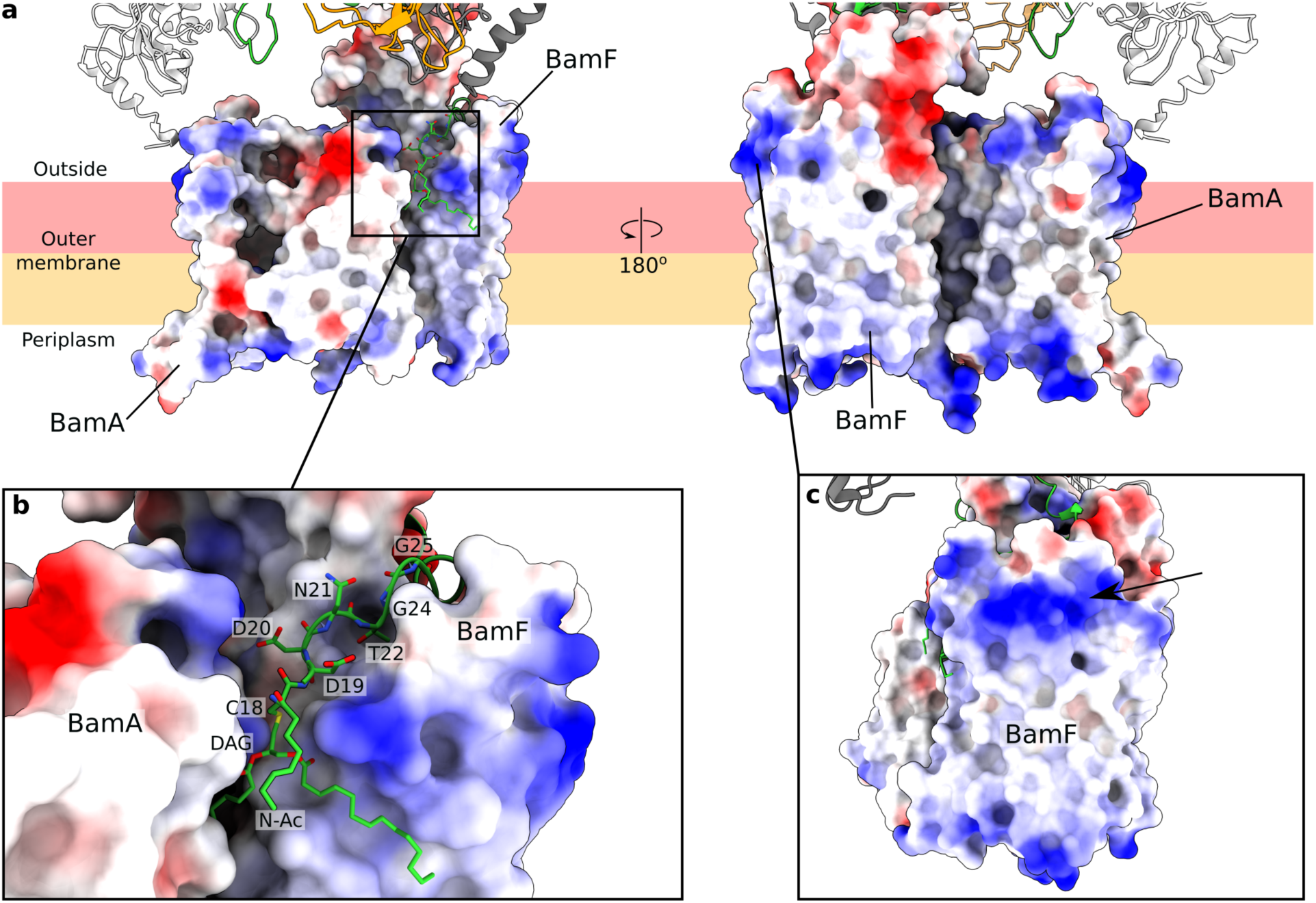
BtBamAF surface charges. **a**, Surface electrostatics representation of BtBamA and BtBamF, with positive charge in blue and negative charge in red. **b**, Close-up view of the BamG lipid anchor and LES (green), which interacts with positive patches formed by BamF and BamA extracellular loops. DAG, diacylglycerol; N-Ac, N-acetyl chain. **c**, A 90° rotated view of BamF compared to panel (**a)** shows a positive patch at the outside OM interface (arrow) that is facing away from the rest of the BAM complex.

**Extended Data Figure 10.**
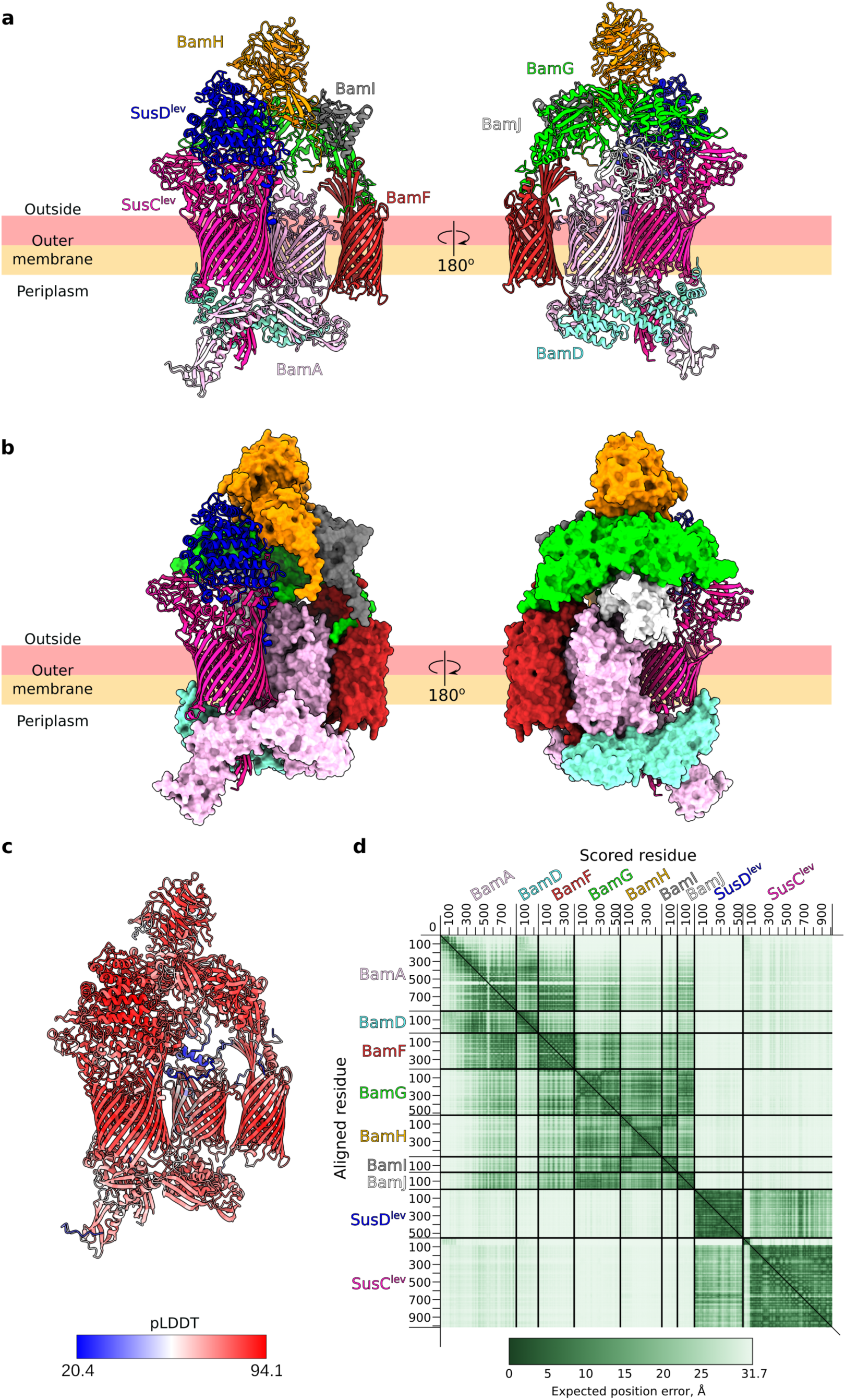
AlphaFold3 prediction of a BtBAM-SusCD^lev^ complex. The top-scoring AF3^102^ prediction of BtBAM with the levan transporter complex SusCD^lev^, shown in cartoon representation (**a**) and with the BtBAM components shown as solvent-accessible surfaces (**b**). The ipTM and pTM values for the predicted model were 0.49 and 0.53, respectively. **c**, The BtBAM-SusCD^lev^ complex coloured according to per-residue local confidence. pLDDT, predicted local distance difference test. **d**, Predicted aligned error plot for the BtBAM-SusCD^lev^ complex. The plot was visualised using PAE Viewer^103^.

