## Supplementary Information for "Discovery of a distinct BAM complex in the Bacteroidetes"


Augustinas Silale[#1], Mariusz Madej[#2*], Katarzyna Mikruta[2,3], Andrew M. Frey[1], Adam J. Hart[1], Arnaud Baslé[1], Carsten Scavenius[4], Jan J. Enghild[4], Matthias Trost[1], Robert P. Hirt[1] and Bert van den Berg[1*]

[#]Authors contributed equally to this work

[1]Biosciences Institute, Faculty of Medical Sciences, Newcastle University, Newcastle upon Tyne, NE2 4HH, UK

[2]Department of Microbiology, Faculty of Biochemistry, Biophysics and Biotechnology, Jagiellonian University, 30-387 Krakow, Poland

[3]Doctoral School of Exact and Natural Sciences, Jagiellonian University, 30-348 Krakow, Poland

[4]Interdisciplinary Nanoscience Center (iNANO), Aarhus University, 8000 Aarhus C, Denmark.

* To whom correspondence should be addressed.

### Supplementary Figures

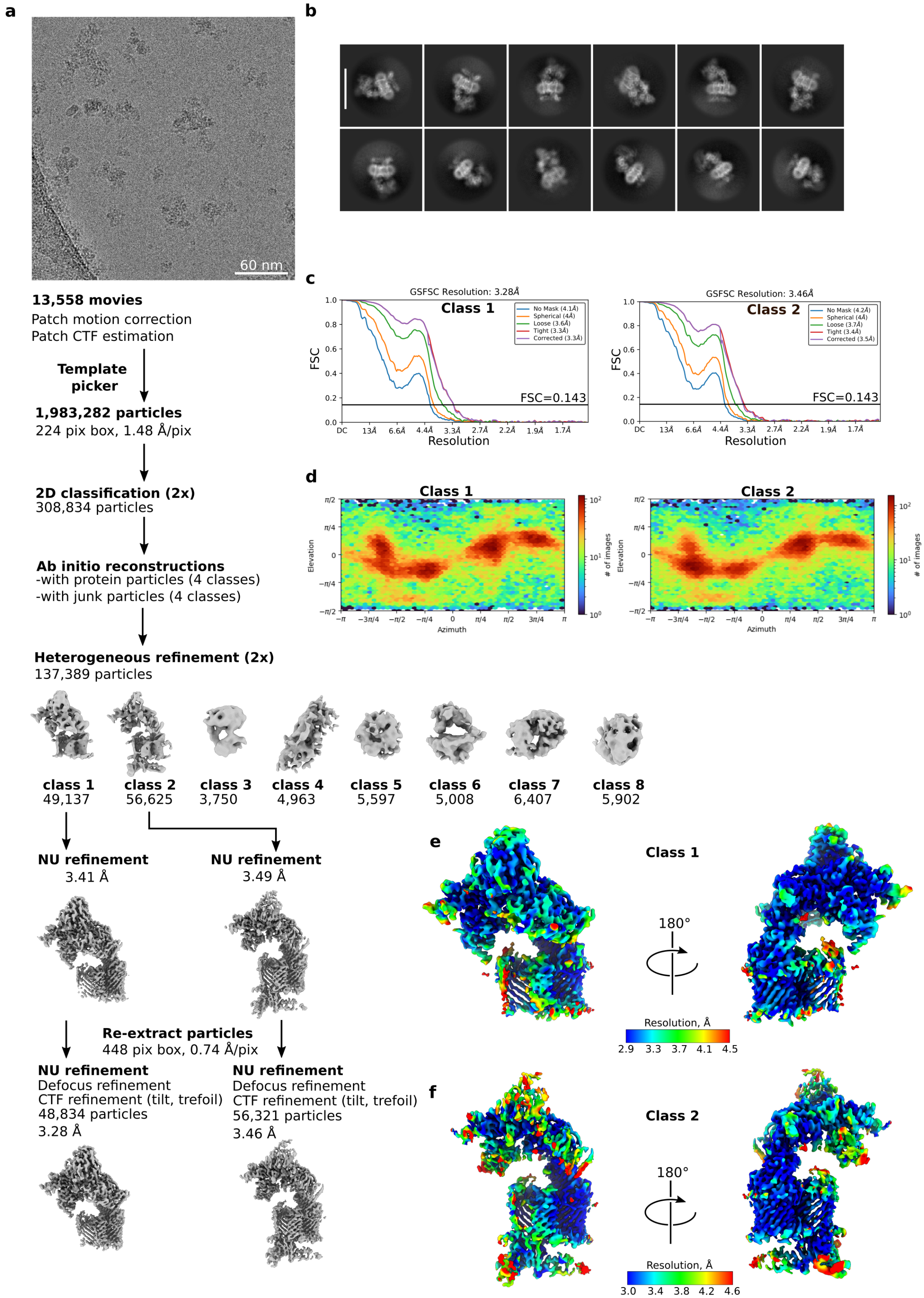

**Supplementary Figure 1. BtBAM cryo-EM data processing workflow.** **a**, Representative motion-corrected movie (n=13,558) and cryoSPARC data processing workflow. **b**, Representative 2D class averages. The white bar represents a length of ~165 Å. **c**, Global gold-standard Fourier shell correlation (FSC) curves. **d**, Viewing direction distribution plots. **e** and **f**, Local resolution estimation for the final class 1 and class 2 maps, respectively

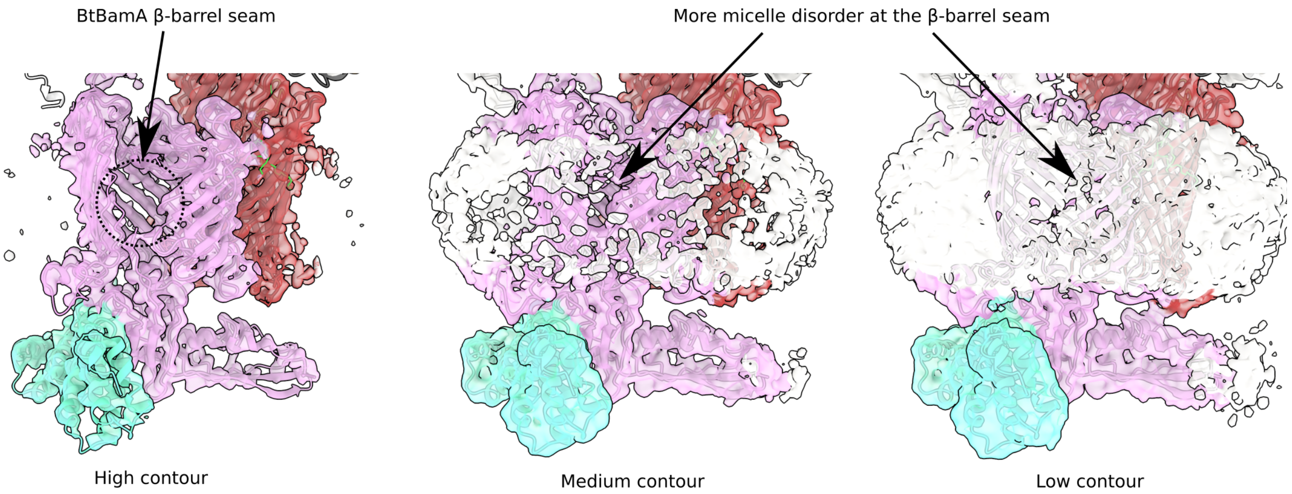


**Supplementary Figure 2. Increased disorder at the BtBamA β-barrel seam**. The BtBamAFD model (cartoon) overlaid with the cryo-EM class 2 map (transparent surface) is displayed at three different contour levels: high, medium and low. As the threshold is lowered, the detergent micelle density does not appear uniformly around the BamA β-barrels. The density is weaker at the β-barrel seam, suggesting that there is more disorder in this region.

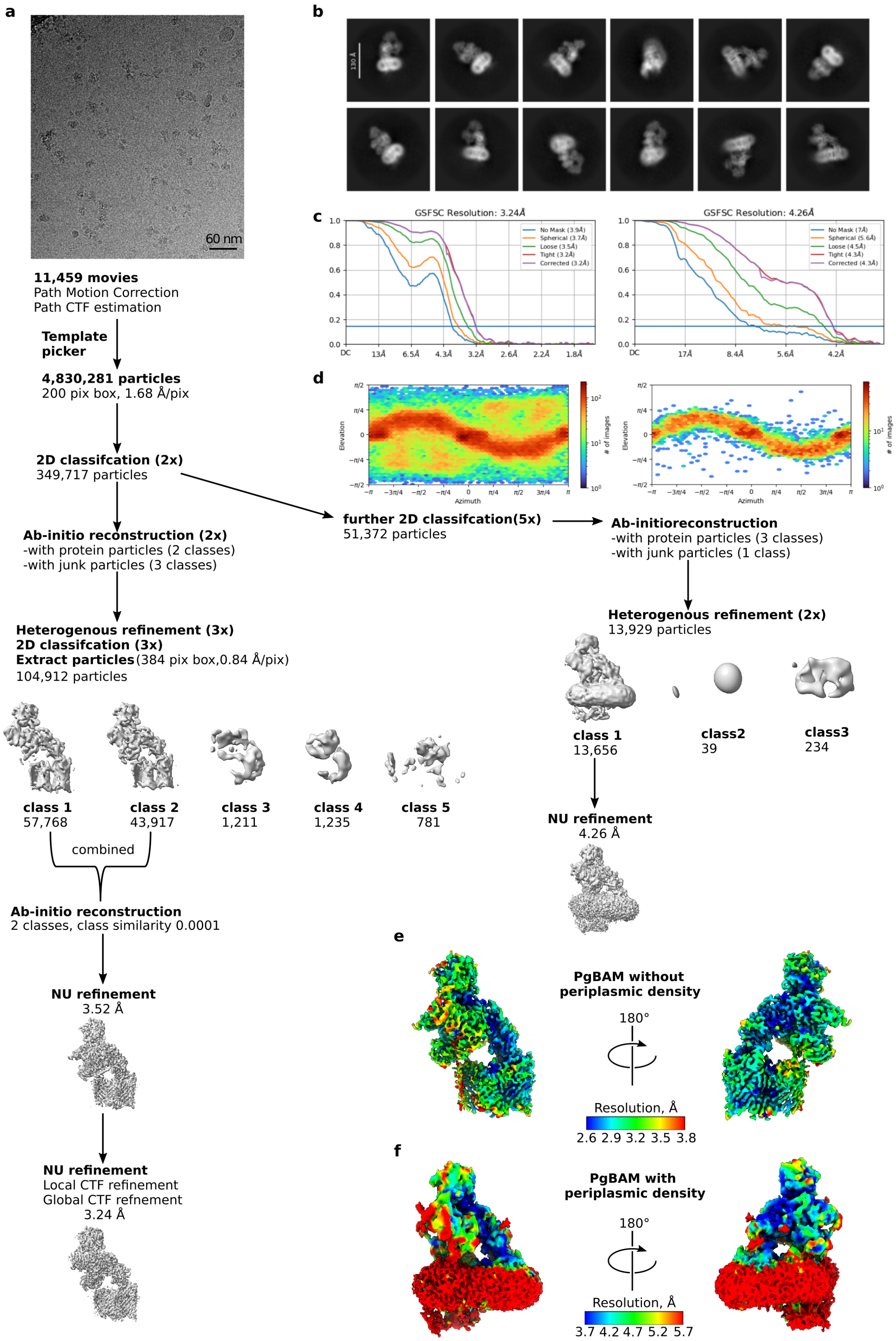
a
60 nm
11,459 movies
Path Motion Correction
Path CTF estimation
Template picker
4,830,281 particles
200 pix box, 1.68 Å/pix
2D classifcation (2x)
349,717 particles
Ab-initio reconstruction (2x)
-with protein particles (2 classes)
-with junk particles (3 classes)
Heterogenous refinement (3x)
2D classifcation (3x)
Extract particles (384 pix box,0.84 Å/pix)
104,912 particles
class 1
57,768
class 2
43,917
class 3
1,211
class 4
1,235
class 5
781
combined
Ab-initio reconstruction
2 classes, class similarity 0.0001
NU refinement
3.52 Å
NU refinement
Local CTF refinement
Global CTF refnement
3.24 Å
b
130 Å
c
GSFSC Resolution: 3.24Å
No Mask (3.9Å)
Spherical (3.7Å)
Loose (3.5Å)
Tight (3.2Å)
Corrected (3.2Å)
GSFSC Resolution: 4.26Å
No Mask (7Å)
Spherical (5.6Å)
Loose (4.5Å)
Tight (4.3Å)
Corrected (4.3Å)
d
Elevation
Azimuth
### of images
further 2D classifcation(5x)
51,372 particles
Ab-initioreconstruction
-with protein particles (3 classes)
-with junk particles (1 class)
Heterogenous refinement (2x)
13,929 particles
class 1
13,656
class2
39
class3
234
NU refinement
4.26 Å
e
PgBAM without
periplasmic density
180°
Resolution, Å
2.6 2.9 3.2 3.5 3.8
f
PgBAM with
periplasmic density
180°
Resolution, Å
3.7 4.2 4.7 5.2 5.7

**Supplementary Figure 3. PgBAM cryo-EM data processing workflow.** **a**, Representative motion-corrected movie (n=11,459) and cryoSPARC data processing workflow. **b**, Representative 2D class averages. The white bar represents a length of 130 Å. **c**, Global gold-standard Fourier shell correlation (FSC) curves. **d**, Viewing direction distribution plots. **e** and **f**, Local resolution estimation for the final reconstructions without and with periplasmic density, respectively.

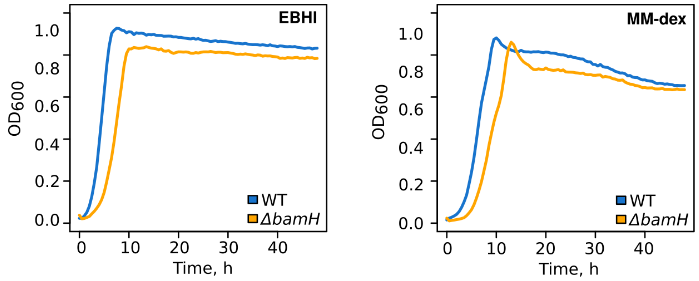


**Supplementary Figure 4. Growth curves of *B. theta* Δ*bamH* strain.** Cells were cultured either in EBHI or in minimal medium supplemented with 0.4% dextran 40 and 10% BHI (MM-dex). Each trace is an average of n=3 technical repeats.

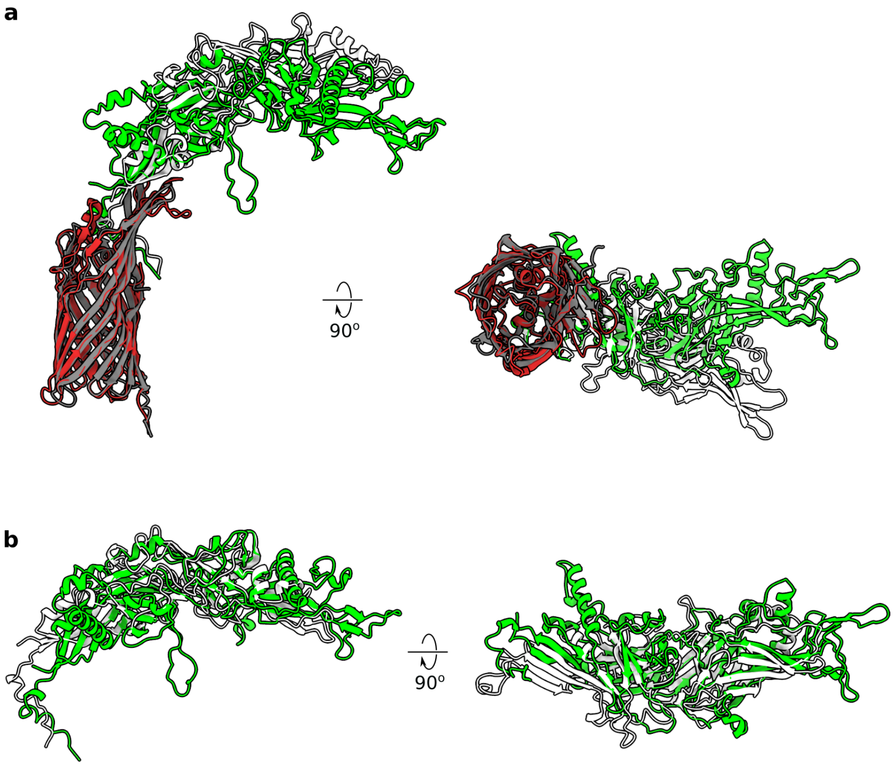


**Supplementary Figure 5. Structural comparison of BtBamFG and Bt1785-86. a**, Superposition of the BtBamFG cryo-EM structure (red, green) and AF3[1] prediction of the Bt1785-86 complex (grey, white). The superposition was generated using Matchmaker in ChimeraX[2] on BtBamF and Bt1785; Cα-Cα RMSD was 1.01 Å between 249 pruned atom pairs. **b**, Superposition of the BtBamG cryo-EM structure and the Bt1786 AF3 prediction. The superposition was generated using Matchmaker in ChimeraX on BtBamG and Bt1786; Cα-Cα RMSD was 1.25 Å between 67 pruned atom pairs, and 15.41 Å between all 385 pairs.

#### Supplementary Movie

**Supplementary Movie 1.** The movie shows a trajectory starting from the PgBAM conformation, morphing to the BtBAM conformation, and back to the PgBAM conformation. The trajectory was generated using the ‘morph’ command in ChimeraX. The protein models are displayed as coloured solvent-accessible surfaces.

#### Supplementary Tables

**Supplementary Table 1.** Cryo-EM data collection, processing and model refinement parameters.

| | BtBAM | | | PgBAM | |
|---|---|---|---|---|---|
| **Data collection** | | | | | |
| Electron microscope | Titan Krios | | | Titan Krios | |
| Voltage (kV) | 300 | | | 300 | |
| Spherical aberration (μm) | 2.7 | | | 2.7 | |
| Camera | Falcon 4i (counting) | | | Gatan K3 | |
| Energy filter | Selectris (10 eV slit) | | | BioQuantum (20 eV slit) | |
| Magnification | 165,000x | | | 105,000x | |
| Pixel size (Å) | 0.74 | | | 0.84 | |
| Total dose ($e^-$/Å$^2$) | 35 | | | 41 | |
| Defocus range (μm) | -0.8 to -2.0 | | | -0.6 to 1.5 | |
| Number of movies collected | 13,558 | | | 11,459 | |
| **Image Processing** | **Class 1** | **Class 2** | | **Class 1** | **Class 2** |
| Symmetry | C1 | C1 | | C1 | C1 |
| Initial number of particles | 2,266,553 | 2,266,553 | | 4,830,281 | 4,830,281 |
| Final number of particles | 48,834 | 56,321 | | 86,584 | 13,929 |
| Global resolution (FSC = 0.143) | 3.28 Å | 3.46 Å | | 3.24 | 4.26 |
| Map sharpening B-factor (Å$^2$) | -58.0 | -64.9 | | -82.6 | -49.3 |
| **Refinement** | **BtBamGHIJ** | **BtBamADF** | **Complete BtBAM** | **PgBAM** | **Class 2** |
| Model composition | | | | | |
| Non-hydrogen atoms | 10,517 | 9,004 | 19,521 | 15,298 | - |
| Protein residues | 1,325 | 1,119 | 2,444 | 1,917 | - |
| R.m.s. deviations | | | | | |
| Bonds lengths (Å) | 0.004 | 0.005 | 0.006 | 0.005 | - |
| Bond angles (˚) | 0.792 | 0.964 | 0.934 | 0.785 | - |
| Validation | | | | | |
| MolProbity score | 1.48 | 1.96 | 1.99 | 2.05 | - |
| Clash score | 2.70 | 11.45 | 11.21 | 10.48 | - |
| Rotamer outliers (%) | 0 | 0 | 0 | 0 | - |
| Ramachandran plot | | | | | |
| Favoured (%) | 93.52 | 94.31 | 93.51 | 91.74 | - |
| Outliers (%) | 0.08 | 0 | 0.04 | 0.21 | - |
| PDB | 9HIS | 9HIV | 9HJ3 | 9HJM | - |
| EMDB | EMD-52200 | EMD-52202 | EMD-52209 | EMD-52218 | EMD-52219 |

**Supplementary Table 2.** BlastP search results using BamF sequence as query against the NCBI RefSeq database.

**Supplementary Table 3.** *B. theta bamG* deletion *vs* wild type total membrane fraction proteomics results shown as protein abundance differences.

**Supplementary Table 4.** *P. gingivalis bamG* deletion *vs* wild type whole cell proteomics results shown as protein abundance differences.

**Supplementary Table 5.** Strains used in this study.

| Strain | Relevant genotype | Source |
|---|---|---|
| ***E. coli*** | | |
| **TOP10** | F- *mcrA* Δ(*mrr-hsd*RMS-*mcr*BC) Φ80*lac*ZΔM15 Δ*lac*X74 *rec*A1 *ara*D139 Δ(*araleu*)7697 *gal*U *gal*K *rps*L (StrR) *end*A1 *nup*G | Invitrogen |
| **S17-1λpir** | *TpR SmR recA thi pro hsdR-M+RP4:2-Tc:Mu:Km* Tn7 *λpir.* | ATCC |
| ***P. gingivalis*** | | |
| **ATCC33277** | Wild type | [3] |
| **RagB-8His ATCC33277** | *ragB* p.$N^{503}$_extHHHHHHHH ($Tet^R$) | [4] |
| ***ΔbamG* ATCC33277** | *ΔbamG* (NCBI:PGN_1735)($Tet^R$) | This study |
| ***ΔbamH* ATCC33277** | *ΔbamH* (NCBI:PGN_0296)($Tet^R$) | This study |
| ***ΔbamJ* ATCC33277** | *ΔbamJ* (NCBI:PGN_1188)($Tet^R$) | This study |
| **BamG-7His ATCC33277** | *bamG* (NCBI:PGN_1735) p.$N^{455}$_extHHHHHHH ($Tet^R$) | This study |
| **RagB-8His in *ΔbamG* ATCC33277** | *ΔbamG ragB* p.$N^{503}$_extHHHHHHHH ($Em^R$)($Tet^R$) | This study |
| ***B. theta*** | | |
| **VPI-5482 *tdk*$^-$** | *tdk*$^-$ | [5] |
| **Bt1761$_{his}$** | *tdk*$^-$ *bt1760*$_{D42A}$ *bt1761*$_{his}$ | [6] |
| **Bt1761$_{his}$ *ΔbamG*** | *tdk*$^-$ *bt1760*$_{D42A}$ *bt1761*$_{his}$ *bt4306*$^-$ | This study |
| **BtBamA$_{his}$** | *tdk*$^-$ *bt3725*$_{his}$ | This study |
| **BtBamA$_{his}$ *ΔbamG*** | *tdk*$^-$ *bt3725*$_{his}$ *bt4306*$^-$ | This study |
| ***ΔbamH*** | *tdk*$^-$ *bt3727*$^-$ | This study |
| **Bt1927-ON** | *tdk*$^-$ *bt1927-ON* | [7] |
| **Bt1927-ON *ΔbamG*** | *tdk*$^-$ *bt1927-ON bt4306*$^-$ | This study |
| **SusA$_{his}$** | *tdk*$^-$ *bt3704*$_{his}$ | J. Abellon-Ruiz (unpublished) |

**Supplementary Table 6.** Plasmids used in this study.

| Plasmids | | |
|---|---|---|
| **Plasmid** | **Relevant features** | **Source** |
| **pUC19** | *E. coli* cloning vector, $Amp^R$ | Thermo Fisher Scientific |
| **pExchange** | Plasmid for *Bacteroides* spp. genome engineering by allelic exchange. | [5] |
| ***ΔbamG*** | Plasmid for deletion of *P. gingivalis bamG* gene, derivative of pUC19 | This study |
| ***ΔbamH*** | Plasmid for deletion of *P. gingivalis bamH* gene, derivative of pUC19 | This study |
| ***ΔbamJ*** | Plasmid for deletion of *P. gingivalis bamJ* gene, derivative of pUC19 | This study |
| **BamGall** | Master plasmid for *P. gingivalis bamG* modifications, derivative of pUC19 | This study |
| **BamG-7His** | Plasmid for insertion of $His_7$-tag at C-terminal of *P. gingivalis* BamG, used for purification of the BAM complex, derivative of BamGall | This study |
| **pExBt3725his** | Plasmid for inserting a His7-tag at the N-terminus of *B. theta* BamA, derivative of pExchange | This study |
| **pExBt4306KO** | Plasmid for deletion of *B. theta bamG* gene, derivative of pExchange | This study |
| **pExBt3727KO** | Plasmid for deletion of *B. theta bamH* gene, derivative of pExchange | This study |
