## Supplementary Discussion for "Discovery of a distinct BAM complex in the Bacteroidetes"

BlastP searches[1,2] against 1448 complete and annotated Bacteroidota genomes (RefSeq database, NCBI[3]) with BT4367 used as query, identified 1640 significant hits across 1334 genomes with one to three significant hits per genome (Supplementary Table 2). All but three of the larger genomes (genome sizes 1.9-10.1 Mb with 1490 – 11768 annotated proteins per genome) encoded at least one significant BlastP hit (E-value range 0 to 2.31E-14). All hits with E-values > 1E-14 had alignments with typically low coverage of the query sequence, with some of these corresponding to rather short sequences. We consider these hits not valid BamF homologs, and they are annotated in yellow in Supplementary Table 2. By contrast, only nine hits with E-values ranging from 2.00E-03 to 2.30E-02 were identified among the 110 smaller genomes from endosymbionts[4] (range of genome sizes: 0.14-1.9 Mb, with 173 – 1456 annotated proteins).

There were three free-living Bacteroidetes genomes without significant BlastP hits: *Bacteroides xylanisolvens* strain BFG-514, *Bergeyella porcorum* strain QD2021 and *Flavobacterium* sp. xlx-214. An additional tBlastn search for these genomes identified one significant hit in the genome of the *Bacteroides xylanisolvens* strain BFG-514, consistent with other seven strains from the same species that encode proteins with significant BlastP hits. For *Bergeyella porcorum* strain QD2021 one hit was obtained confirming the weak BlastP hit (Bpo_0594). Inspection of the sequence suggested a sequencing problem in the C-terminal ~80 residues of the protein. An AlphaFold 3[5] prediction confirmed the sequencing problem but suggested that BPO_0594 is a BamF homolog (Supplementary Discussion Figure 1). This is consistent with the significant BlastP hits observed for the other three strains from the same species (Supplementary Table 2). Interestingly, no hits were identified for *Flavobacterium* sp. xlx-214, in sharp contrast to all other 173 genomes from *Flavobacterium* species/strains, which encode at least one significant BlastP hit (Supplementary Table 2).

The smaller genomes are from endosymbiotic species[4] with only nine total hits characterised by small coverage (range of alignment length 61 to 220) of the Bt4367/BamF query sequence and higher E-values (Supplementary Table 2). A tBlastn search of the largest genome among the endosymbionts Candidatus *Amoebophilus asiaticus* strain 5a2, which was identified in a free-living amoeba[6], did not identify any BamF hit.

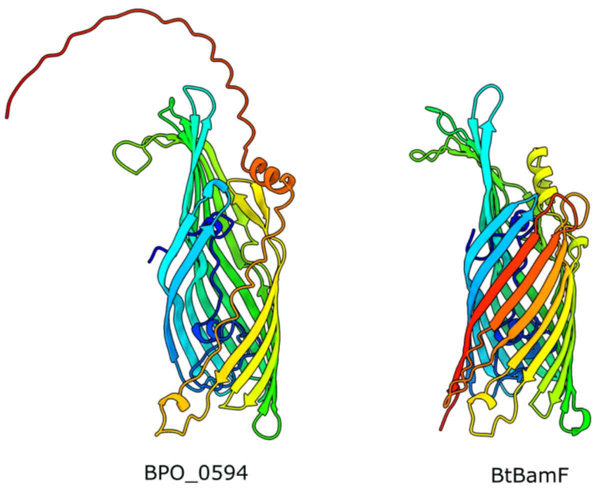


**Supplementary Discussion Figure 1. BPO_0594 structure prediction.** The structure of BPO_0594 (UniProt A0AAU0EZF4) was predicted using AlphaFold 3. BtBamF is shown for comparison; views generated from a superposition. The barrel of BPO_0594 is incomplete, suggesting a possible sequencing or genome assembly error.

### Supplementary Data References
